# Evolution of inflammation and immunity in a dengue virus 1 human infection model

**DOI:** 10.1101/2022.02.10.479935

**Authors:** Adam T. Waickman, Joseph Q. Lu, HengSheng Fang, Mitchell Waldran, Chad Gebo, Jeffrey R. Currier, Liesbeth Van Wesenbeeck, Nathalie Verpoorten, Oliver Lenz, Lotke Tambuyzer, Guillermo Herrera-Taracena, Marnix Van Loock, Timothy P. Endy, Stephen J. Thomas

## Abstract

Dengue virus (DENV) infections are significant sources of morbidity and mortality throughout the tropics and subtropics. Over 400 million infections are estimated to occur every year, resulting in nearly 100 million symptomatic infections and over 20,000 deaths. Early immune response kinetics to infection remain unclear, in large part due to the variable incubation period exhibited by the DENVs after introduction into a susceptible host. To fill this knowledge gap, we performed a comprehensive virologic and immunologic analysis of individuals experimentally infected with the under-attenuated DENV-1 strain 45AZ5. This analysis captured both the kinetics and composition of the innate, humoral, and cellular immune responses elicited by experimental DENV-1 infection, as well as virologic and clinical features. Revealed in this analysis was a robust DENV-specific IgA antibody response that manifested between the appearance of DENV-specific IgM and IgG in all challenged individuals, as well as the presence of a non-neutralizing/NS1-specific antibody response that was delayed relative to the appearance of DENV-virion specific humoral immunity. RNAseq analysis also revealed several distinct and temporally-restricted gene modules that allowed for the identification and differentiation of the innate and adaptive immune response to DENV-infection. Our analysis provides a detailed description, in time and space, of the evolving matrix of DENV-elicited human inflammation and immunity and reveals several previously unappreciated immunological aspects of primary DENV-1 infection that can inform countermeasure development and evaluation.

## INTRODUCTION

Dengue is caused by infection with any of the four dengue-virus (DENV) serotypes (DENV-1 to −4). The viruses are transmitted when an infected female *Aedes* mosquito takes a blood meal from a susceptible, non-immune host [1]. Each year, an estimated 400 million people, most living in tropical and subtropical regions, are thought to be infected, and approximately 100 million infections manifest with a range of clinical phenotypes [2, 3]. Roughly 500,000 dengue cases require hospitalization and up to 20,000 people succumb to the disease every year [4]. Dengue is expected to continue as a worsening global public health challenge because the main drivers of DENV transmission are projected to continue and/or accelerate over the next several decades (e.g., climate change, travel, population growth, poverty, urbanization). It is estimated that by 2080, over 6 billion people will live at daily risk of a DENV infection [5]. While a number of vaccines are in development [6–9], the single dengue vaccine currently available is limited to people 9 years of age and older and only in those who have pre-existing dengue immunity [10–12]. While several promising candidates are in early-stage clinical development [13], there is no anti-DENV antiviral currently available for prophylaxis or as a therapeutic. Therefore, despite more than century of basic science research, countermeasure development efforts, and attempts at vector control, dengue remains a largely unchecked public health burden.

Development of dengue countermeasures (i.e. drugs, vaccines, diagnostics, vector control tools) has been hindered by fundamental gaps in our understanding of DENV transmission, pathogenesis, and anti-DENV immune responses. These knowledge gaps include understanding who is at risk of DENV exposure, why do some infected people become ill and others do not, and what environmental and/or genetic factors increases a person’s infection risk. Addressing these knowledge gaps is critical to support countermeasure development efforts. Studying wild type infections to answer these questions has been challenging for a number of reasons including; 1) it is extremely difficult to capture people in the first few days after infection and before symptoms develop; 2) many people living in dengue endemic regions have pre-existing DENV or non-DENV flavivirus immunity from past infections or vaccinations; and 3) it is difficult to collect blood samples with a frequency which allows for detailed kinetic analyses. For these reasons, our group has collaboratively developed an experimental Dengue Human Infection Model (DHIM) [14].

A DHIM has the potential to be a powerful tool for studying human host responses to DENV infection. Individuals with known pre-existing flavivirus immune profiles are infected with a highly characterized, and under-attenuated, DENV strain manufactured under Good Manufacturing Practices and administered in a controlled clinical environment. The participants are then intensely followed measuring clinical, clinical lab, virologic, and immunologic responses over time. Experimental human infection with the DENVs have been documented since 1902 and have greatly contributed to our foundational understanding of DENV transmission, virology, and immunology [15–25]. More recent studies have characterized the clinical and functional immune profiles following DENV-1 or DENV-2 infection, explored the early transcriptional features of attenuated DENV-2 infection, and described the cellular immune responses following infection [9, 14, 26–28]. However, to date, no study has sought to comprehensively and longitudinally assess and integrate the virologic and immunologic parameters associated with a primary DENV infection in a flavivirus-naïve individual.

To this end, our team set out to thoroughly characterize the clinical, immunologic and virologic features of a primary DENV-1 infection in flavivirus-naïve adults with fine temporal resolution. This analysis captured both the kinetics and composition of the innate, humoral, and cellular immune response elicited by experimental DENV-1 infection, as well as, the virologic and clinical features of infection.

## RESULTS

### Study overview and clinical outcome of DENV-1 infection

This study was conducted to characterize the longitudinal virologic and immunologic profile associated with a primary DENV-1 infection in flavivirus naïve adults. To this end, nine participants (**Supplemental Table 1**) were enrolled to receive a single subcutaneous inoculation of 0.5mL of a 6.5×10^4^ PFU/ml suspension of the under-attenuated DENV-1 strain 45AZ5 [14]. This virus strain originated from a Chinese patient with mild dengue fever living on the island of Nauru (Western Pacific) in 1974, and was subsequently serially passaged in a diploid fetal rhesus lung cell line (FRhL) and mutagenized with 5-azacytidine to facilitate the accumulation of attenuating mutations [29, 30].

Participants in this study were evaluated daily for the first 14 days after inoculation, then every other day until 28 days post virus inoculation with additional visits on days 90 and 180 post infection. All nine enrolled participants completed the study per protocol and were included in the subsequent virologic and immunologic analyses.

Within 7 days after inoculation, 5 of 9 participants reported at least 1 solicited local AE (injection site symptom). The 5 participants reporting solicited local AEs described mild (grade 1) reactions including bruising, discomfort, erythema, and injection site hemorrhage. No related unsolicited local AEs were reported. Within 28 days after inoculation all 9 participants reported at least one solicited systemic AE including headache, rash, fever, eye pain, weakness, and myalgia (**Table 1, Supplemental Table 2**). Most participants (8/9) had at least 1 laboratory abnormality between study days 1 and 181, the majority of which were mild or moderate and all were reported within 28 days after inoculation or 7 days post hospitalization, whichever was later (**Table 1, Supplemental Table 3**). At least 1 severe laboratory abnormality was reported by 3 of the 9 participants. One potentially life-threatening AE was observed (low serum glucose), but failed to correlate with the clinical presentation at the time of the routine follow-up visit and returned to normal levels by the following day. The abnormality was subsequently assessed by the principal investigator as unrelated to the viral challenge. As with previous studies, all participants who became ill were managed with oral fluids, acetaminophen, and an anti-nausea medication if required. No intravenous access was required.

**Table 1.** DHIM proposed performance parameters.

| **Participants experiencing:** | **All Participants**<br>**(N=9) n/M (%)** |
| --- | --- |
| Measurable RNAemia of 3 to 11 days duration | 9/9(100) |
| Fever measured at least 2 times in 24 hours but not lasting more than 72 hours | 5/9(55.6) |
| At least one of the following symptoms | 9/9(100) |
| At least 2 of the following symptoms | 8/9(88.9) |
| Fever and at least one of the following symptoms | 5/9(55.6) |
| Fever and at least 2 of the following symptoms | 4/9(44.4) |
| Headache, maximum grade 1 or 2 | 8/9(88.9) |
| Myalgia, maximum grade 1 or 2 | 7/9(77.8) |
| Rash, maximum grade 1 or 2 | 5/9(55.6) |
| Liver function tests (ALT, AST), maximum grade 1 or 2 | 3/9(33.3) |
| Leukopenia, maximum grade 1 or 2 | 6/9(66.7) |
| Thrombocytopenia, maximum grade 1 or 2 | 1/9(11.1) |

All study participants developed detectable RNAemia and viremia as assessed by qRT-PCR and Vero cell-based plaque assay, respectively, with a largely uniform viral kinetics pattern (**Figure 1A, Figure 1B, Supplemental Figure 1**). The incubation period (time from infection to detectable RNAemia) ranged from 6 to 11 days, with RNAemia persisting for a mean of 8 days. Peak viral load ranged from 7.46 × 10^5^ to 8.33 × 10^7^ genome equivalents (GE)/mL, or 1.3 × 10^4^ to 6.8 × 10^4^ plaque forming units (PFU)/mL of serum. In addition to circulating virus, DENV NS1 antigenemia was detected as early as Day 7 post inoculation (range 7 – 13 days), with NS1 antigenemia persisting for a mean of 13 days across all study participants (**Figure 1C**). When compared to RNAemia as quantified by qRT-PCR, levels of NS1 serum antigenemia exhibited more variability between participants and appeared on average two days later than viral RNA (**Figure 1D**).

**Figure 1.**
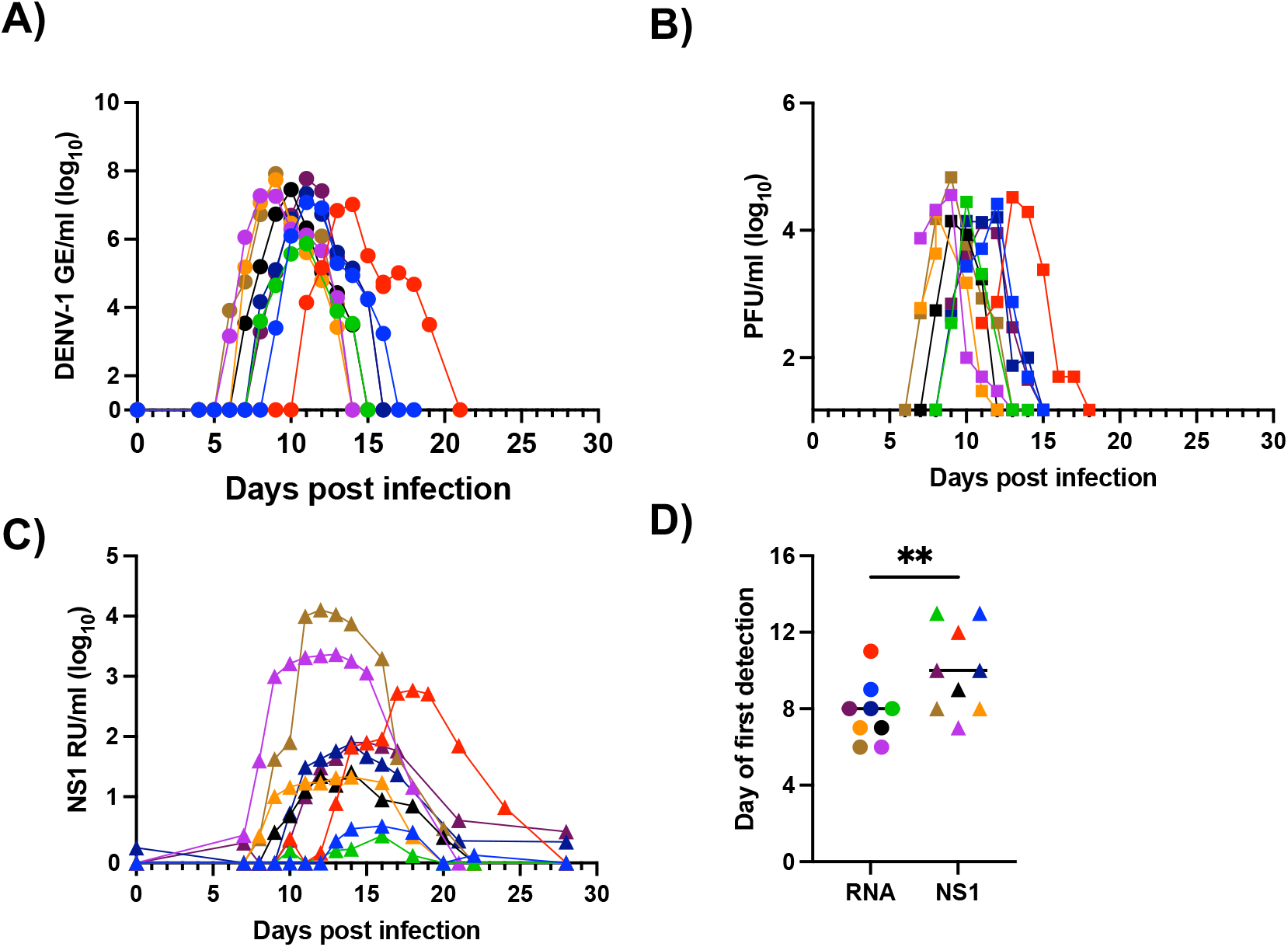
Kinetics of DENV-1 infection in 45AZ5 challenged individuals. **A**) DENV-1 RNA content in serum as assessed by qRT-PCR. **B**) Infectious DENV-1 content in serum as assessed Vero cell plaque assay. **C**) NS1 protein content in serum as assessed by ELISA. **D**) Comparison of RNAemia and NS1 antigenemia onset in all study participants. ** < 0.01 paired T test. Line indicates group median.

### Kinetics and specificity of DENV-1 elicited humoral immunity

Having established the kinetics of DENV-1 RNAemia, viremia, and antigenemia following 45AZ5 infection, we next assessed the induction of DENV-1 specific humoral immunity. While induction of IgM or IgG isotype antibodies is most commonly used for serological confirmation of DENV infection and for monitoring durable antiviral immunity, we included IgA in our analysis given our previous observation that primary DENV infection is associated with a significant IgA response and may have an association with disease severity [31, 32]. Using a previously described virion-capture ELISA assay we observed a robust - though transient - DENV-1 specific IgM and IgA response in all participants, with IgM and IgA seroconversion occurring on average by post-infection day 14 and 16, respectively (**Figure 2A, 2B**). All participants still exhibited detectible anti-DENV-1 IgM titers at day 90 post DENV-1 challenge, while only 3/9 participants maintained an IgA titer. DENV-1 specific IgG seroconversion occurred later than IgM or IgA in all participants, with detectable DENV-1 specific IgG appearing on average by day 19 post infection and remaining stable out to 90 days post infection (**Figure 2A, 2B**).

**Figure 2.**
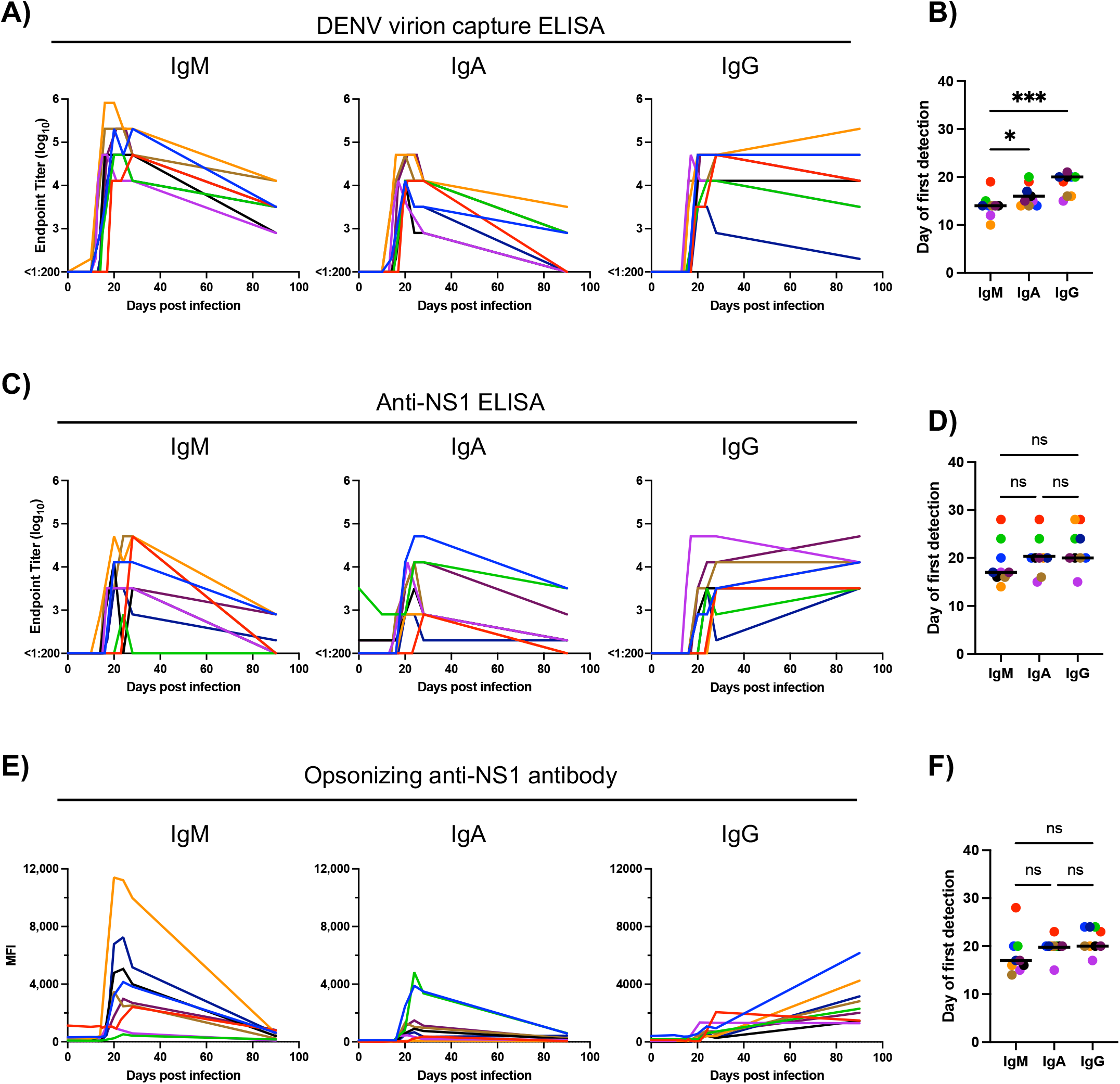
Kinetics and characteristics of DENV-1 specific humoral immunity following 45AZ5 infection. **A**) Antibody endpoint titers using virion-capture ELISA. **B**) Day of DENV-virion specific seroconversion by antibody isotype. * < 0.05, ** < 0.01 paired one-way ANOVA with correction for multiple comparisons. **C**) Antibody endpoint titers using NS1 protein ELISA. **D**) Day of anti-NS1 seroconversion by antibody isotype. **E**) Opsonizing anti-NS1 antibody staining using NS1 expressing CEM.NK^R^ cells. **F**) Day of opsonizing anti-NS1 seroconversion by antibody isotype

In light of the robust NS1 antigenemia observed following 45AZ5 infection, we additionally assessed the kinetics and magnitude of anti-NS1 specific humoral immunity. Similar to what was observed with the DENV-1 virion capture assay, DENV1-NS1 specific IgM/IgA responses following 45AZ5 infection were transient, while IgG levels persisted and remained elevated out to day 90 post infection (**Figure 2C**). DENV-1-NS1 specific IgM/IgA response occurred earlier (on average on Days 19/20, respectively) when compared to IgG responses which appeared, on average, by Day 23 (**Figure 2D**). When comparing DENV-virion-specific with DENV-1-NS1-specific humoral immune response, anti-virion immunity tended to appear earlier (average seroconversion was between Day 14 to 19 for the anti-virion response versus Day 19 to 23 for the anti-NS1 response) (**Figure 2B and 2D**), similar to the pattern observed for the appearance of DENV-1 virions and NS1 antigen.

NS1 is predominantly secreted as a hexamer by flavivirus infected cells, but the protein can also be found on the cellular plasma membrane in a dimeric configuration [33, 34]. While the biological function of surface-restricted NS1 is unclear, the antigen can facilitate opsonization of infected cells by NS1-reactive antibodies, thereby allowing NK cells, monocytes and other phagocytic cells to recognize and clear infected cells via Fc receptor mediated mechanisms such as antibody-dependent cellular cytotoxicity (ADCC) and antibody-dependent cellular phagocytosis (ADCP) [35, 36]. To measure the abundance of opsonizing NS-1 specific antibodies we utilized a CEM.NKR cell line that was engineered to stably express DENV-1 NS1 (**Supplemental Figure 2**) and quantified the abundance of anti-NS1 opsonizing antibodies using flow cytometry. As was observed for the anti-hexametric NS1 antibody titers, induction of opsonizing anti-NS1 IgM and IgA isotype antibodies was transient after inoculation, with IgG isotype responses appearing later and persisting through day 90 post infection (**Figure 2E**). The average day of opsonizing anti-NS1 seroconversion was Day 18 (IgM), Day 20 (IgA) and Day 21 (IgG) (**Figure 2F**), further emphasizing the delayed kinetics of anti-NS1 immunity relative to virion-specific humoral immunity.

### Kinetics and specificity of DENV-1 elicited cellular immunity

To define the magnitude and specificity of DENV-1 elicited cellular immunity following 45AZ5 infection, PBMC collected from all study participants on study days 0, 28, and 90 were stimulated with overlapping peptide pools spanning the E, NS1, NS3, and NS5 proteins of DENV-1 and analyzed in an IFN-γ ELISPOT assay (**Supplemental Table 4**). As has been described following natural DENV infection or inoculation with a Live Attenuated Vaccine (LAV) product [37, 38], infection with 45AZ5 resulted in the generation of significant DENV-1 specific cellular immunity - as assessed by IFN-γ ELISPOT – by 28 days post infection that remained elevated out to day 90 post infection (**Figure 3A, Supplemental Figure 3**). DENV-1 NS3 was the dominant antigen recognized on day 28 as well as day 90 post infection (**Figure 3B, 3C, Supplemental Figure 3**), followed in magnitude by E, NS3 and NS1.

**Figure 3.**
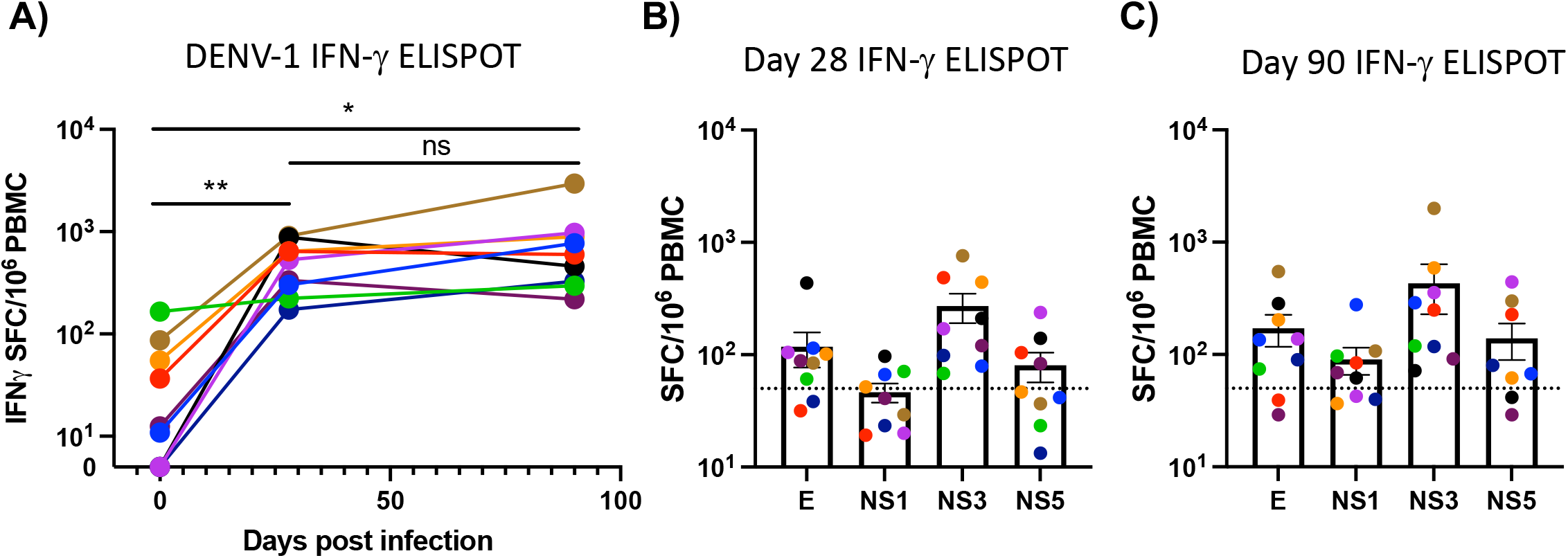
Kinetics and specificity of DENV-1 elicited cellular immunity. **A**) Total DENV-1 specific cellular immunity as assessed by IFN-γ ELISPOT. Sum of background-subtracted E, NS1, NS3, and NS5 reactivity at each indicated time point. * < 0.05, ** < 0.01 paired one-way ANOVA with correction for multiple comparisons. **B**) Antigen specific breakdown of day 28 DENV-1 specific cellular immunity. **C**) Antigen specific breakdown of day 90 DENV-1 specific cellular immunity

### Transcriptional analysis of DENV-elicited inflammation

In light of the dramatic serologic and cellular immune responses elicited by 45AZ5 infection, we next endeavored to better define the early kinetic profile of DENV-1 elicited inflammation. Previous studies have examined the transcriptional profile associated with acute natural DENV infection and have identified conserved gene signatures that correlate with disease severity [39]. However, the inability of these studies to precisely define the day of DENV infection means that the timing and nature of the earliest transcriptional response to DENV infection is unclear.

To address this, we performed RNAseq analysis on whole blood collected on study days 0, 8, 10, 14, and 28 from all 9 study participants. These study days were selected based on previously scRNAseq analysis of a select number of participants enrolled in a separate DENV-1 experimental infection study which demonstrated that these days would likely capture both the early innate/inflammatory signatures associated with acute DENV infection as well as the subsequent activation and expansion of DENV-reactive lymphocytes [26].

Consistent with our previously published results, only modest and inconsistent transcriptional perturbations from baseline were observed across study participants on day 8 post infection. However, significant transcriptional shifts away from pre-infection profiles were observed on days 10 and 14 post infection, with all participants demonstrating a return to baseline by day 28 post infection (**Figure 4A**). No statistically-significant differentially expressed genes (DEGs) were observed on days 8 and 28 relative to day 0. However, 112 genes (110 upregulated, 2 downregulated) were observed to be differentially expressed on day 10 post infection (**Figure 4B, Supplemental Table 5, Supplemental Table 6**), while 177 genes (151 upregulated, 26 downregulated) were differentially expressed on day 14 relative to pre-infection (**Figure 4C, Supplemental Table 7, Supplemental Table 8**).

**Figure 4.**
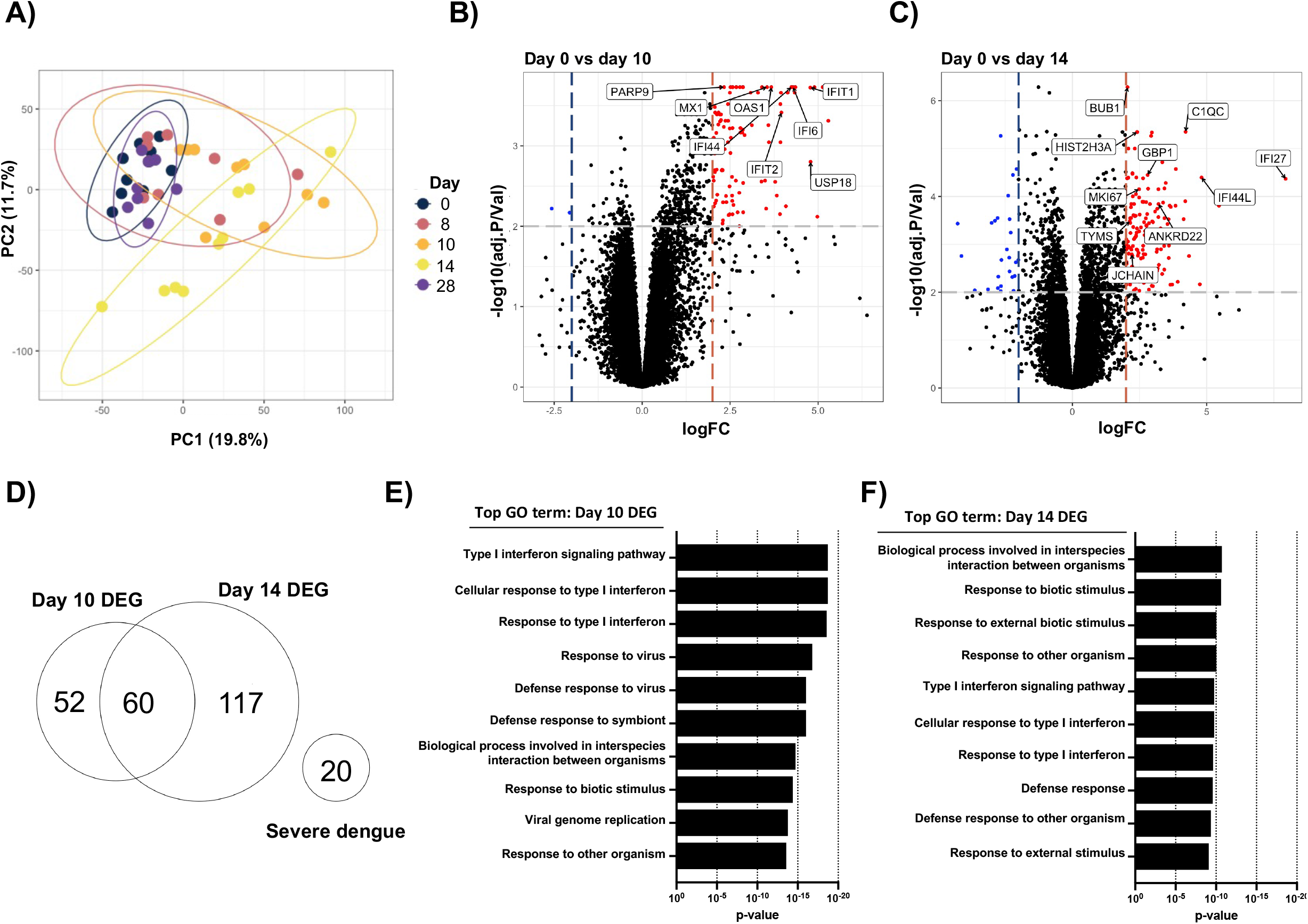
Kinetics and composition of DENV-1 elicited inflammation. **A**) PCA plot of RNAseq analysis of whole blood obtained on days 0, 8, 10, 14, and 28 post DENV-1 infection. Points colored by sample collection day. **B**) Volcano plot showing differential gene expression day 0 vs day 10 post infection with select statically and biologically significant genes highlighted. Genes with a log_2_ fold change of >2 and an adjusted p-value < 0.01 were considered significant. **C**) Volcano plot showing differential gene expression day 0 vs day 14 post infection with select statically and biologically significant genes highlighted. Genes with a log_2_ fold change of >2 and an adjusted p-value < 0.01 were considered significant. **D**) Overlap of Differentially Expressed Genes (DEG) observed on days 10 and 14 and relationship to genes previously described to correlate with a progression to severe dengue. **E**) Gene Ontogeny (GO) analysis of gene differentially expressed on day 10 relative to day 0 post DENV-1 infection. **F**) GO analysis of genes differentially expression day 14 relative to day 0 post DENV-1 infection

The DEGs observed on day 10 post infection included many canonical interferon-stimulated and antiviral gene products (MX1, IFIT1, PARP9, IFI44), while the day 14 DEG additionally included genes associated with lymphocyte proliferation and antibody production and functionalization (MKI67, TYMS, JCHAIN, C1QC). There was significant overlap between the DEG observed on days 10 and 14 post 45AZ5 infection, mostly restricted to gene products associated with Type I interferon signaling and response to viral ligands (**Supplemental Table 9**). Consistent with the clinical profile associated with 45AZ5 experimental infection, there was no overlap between the DEGs capture in this study and a previously published gene set curated to identify participants with an increased risk of progressing to severe dengue [39] (**Figure 4D**). Gene Ontogeny (GO) analysis performed on the DEGs identified on days 10 and 14 post infection indicated that the majority of the gene products differentially expressed on day 10 post 45AZ5 infection corresponded to pathways associated with acute/innate immune responses to cytokine stimulation and/or viral antigens (**Figure 4E**), while those genes differentially expressed on day 14 post infection also included terms associated with lymphocyte activation and the induction of an adaptive immune response (**Figure 4F**).

#### Kinetics DENV-elicited inflammation and lymphocyte activation

Having identified DEG on days 10 and 14 following DENV-1 infection, we next endeavored to better describe the functional kinetics of DENV-1 elicited inflammation and immunity using these transcriptional profiles. The day 14 DEG contained gene products that corresponded to both Type I IFN signaling and virus sensing (IFI27, IFI44L, GBP1), as well as genes associated with lymphocyte activation and proliferation (MKI67, TYMS, JCHAIN, MZB1).

Leveraging this functional heterogeneity, we grouped the day 14 DEG into statistically distinct gene modules that exhibited conserved patterns of expression across all time points captured in this analysis. Using this approach, 3 distinct gene modules were identified within the aggregated dataset that exhibited coordinated patterns expression following DENV-1 infection (**Figure 5A**). Gene Module 1 identified in this analysis consisted of gene products associated with cellular proliferation, antibody secretion, and functional antibody responses (**Figure 5A, Figure 5B, Supplemental Table 10**), while Gene Module 2 preferentially contained genes associated with Type I/II Interferon signaling, virus sensing, and defense responses to viruses (**Figure 5A, Figure 5C, Supplemental Table 10**). Gene Module 3 consisted of genes that were suppressed on day 14 relative to day 0 post infection but do not appear to fall into any biologically relevant category (**Supplemental Table 10**).

**Figure 5.**
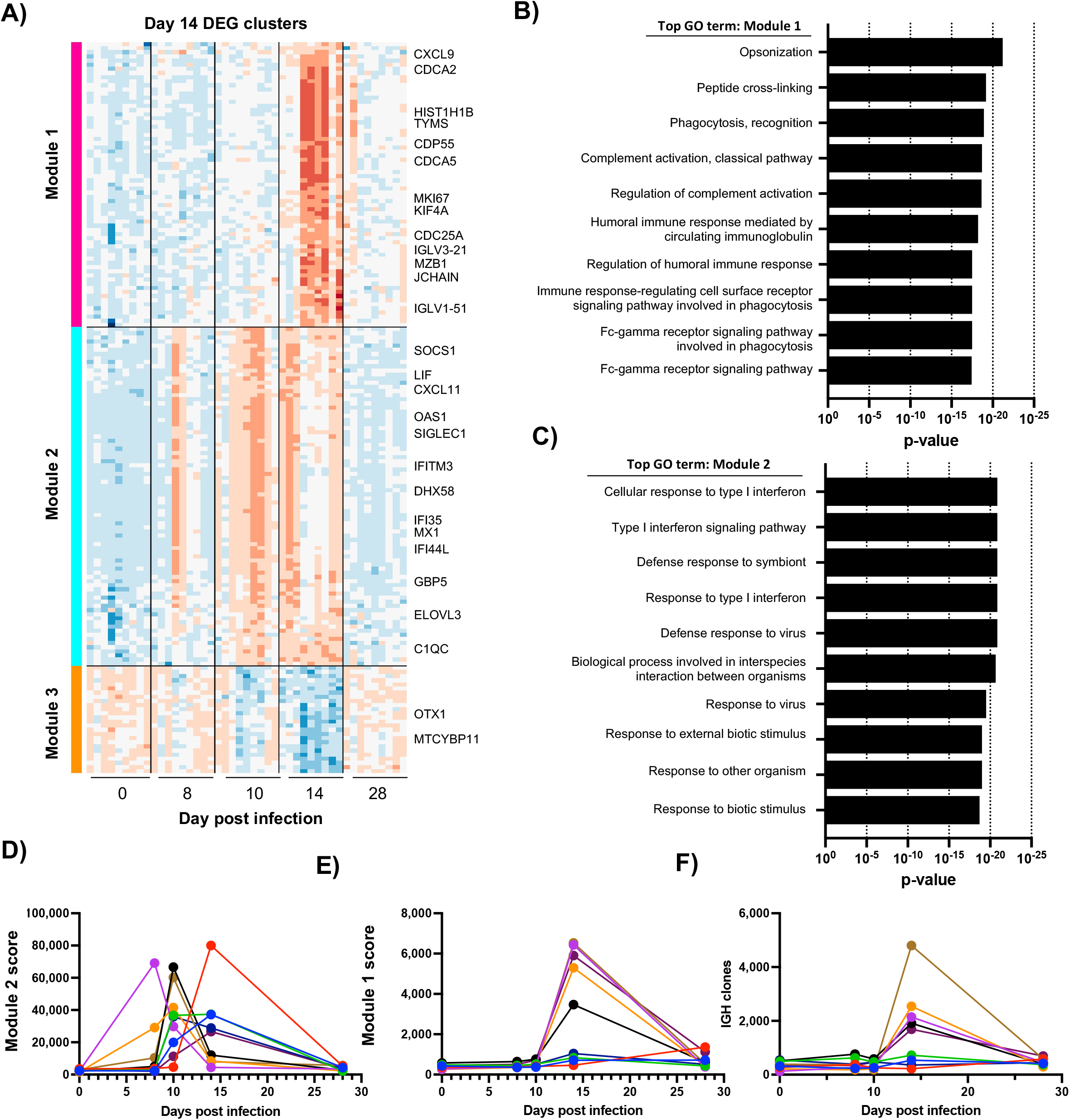
Identification, characterization, and timing of DENV-1 elicited inflammation and lymphocyte activation. **A**) Identification of coordinated gene modules among genes differentially express on day 14 vs day 0 with select biologically significant genes highlighted. **B**) Gene ontology (GO) analysis of the genes found in “Module 1”, exhibiting coordinated upregulation on day 14 relative to day 0. **C**) Gene ontology (GO) analysis of the genes found in “Module 2”, exhibiting coordinated upregulation on days 8, 10, and 14 relative to day 0 post DENV-1 infection. **D**) Kinetics of module 2 expression across all 9 study participants and 5 time points. **E**) Kinetics of module 1 expression across all 9 study participants and 5 time points. **F**) Identification and quantification of unique IGH clones across all 9 study participants and 5 time points from the unenriched RNAseq data using MiXCR.

The expression of genes found in Module 1 or Module 2 were mutually exclusive across the time points captured in this analysis, with the expression of the inflammation-associated Module 2 preceding the expression of the lymphocyte–associated Module 1 in all participants (**Figure 5D, Figure 5E**). The preferential expression of Module 2 genes on day 8 and 10 post infection corresponds to the window of viremia observed in this study, while the expression of Module 1 genes reflects the anticipated timing of B cell activation following primary DENV-1 infection.

To further confirm that Module 1 expression corresponds to B cell activation, we leveraged the fact that activated/antibody-secreting B cells contain ~1,000x the number of immunoglobulin transcripts as resting naïve/memory B cells. We assembled and annotated the unique IGH clones contained within our unenriched/untargeted RNAseq libraries. This analysis indicated that the number of unique antibody clones captured in our data was significantly elevated on day 14 relative to all other time points (**Figure 5F**). Of note, not all participants exhibited a Module 1 and IGH clone peak on day 14, suggesting that these participants either did not experience a significant adaptive response to DENV-1 infection or that the peak of lymphocyte activation occurred outside of our analysis window. In their totality, these data demonstrate the presence of temporally distinct and dynamic gene modules that evolve following DENV-1 infection, and can be used to infer the magnitude of inflammation and timing post DENV-1 infection.

### Evolution of DENV-elicited inflammation/immunity and correlation between virologic and immunologic features of DENV-1 infection

To add context to the virologic and immunologic parameter captured in this analysis, and to explore potential causal relationships with clinically relevant endpoints, we next endeavored to clarify the relative timing and correlative relationships between the individual components captured in this analysis.

We selected 11 virologic and immunologic parameters and calculated the probability of observing a positive signal for a given feature within a 4–5-day window between 0- and 30-days post DENV-1 infection (**Figure 6A**). Placed in context, the delayed appearance of NS1 antigen relative to DENV viral RNA is mirrored by a corresponding delay in the appearance of NS1-specific antibodies relative to virion-specific antibodies. However, a conserved progression of IgM, IgA, then IgG seroconversion is observed for antibodies targeting both viral antigens. The probability of observing the inflammation-associated RNAseq-derived Gene Module 2 coincides with the appearance of viral RNA and is immediately followed by appearance of the lymphocyte-activation associated Gene Module 1 - the time of DENV virion-specific seroconversion. Notably, the inflammation-associated Gene-module 2 can be detected before the window during which fever was observed in study participants.

**Figure 6.**
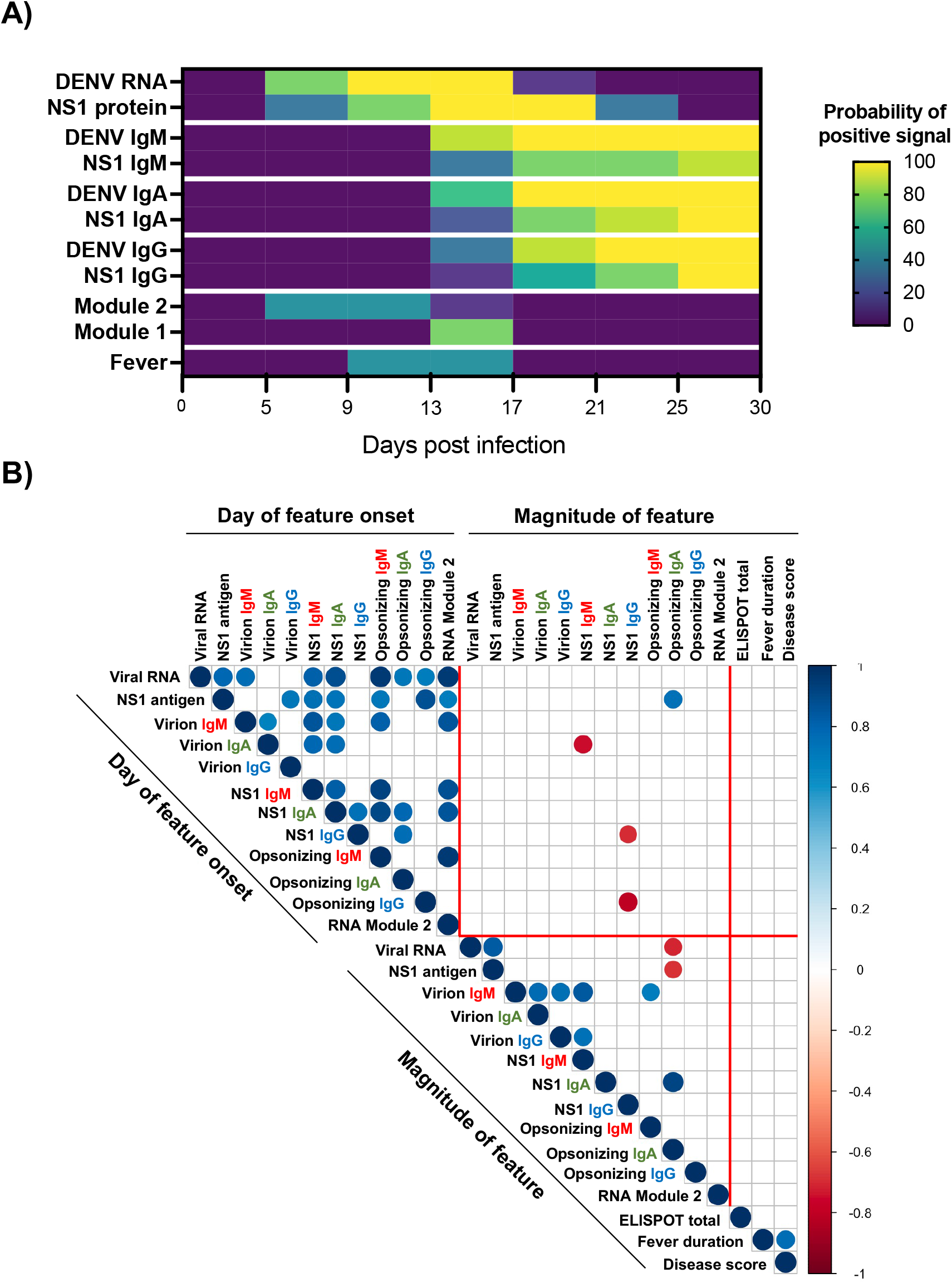
Summary and correlation of DENV-elicited inflammation and immunity. **A**) Aggregate summary of the timing of DENV proliferation and immune response. **B**) Correlation of all virologic and immunologic parameters quantified in this study. Point size and color indicates strength and direction of correlation.

The day of RNAemia onset correlated significantly with the appearance of NS1 antigenemia and the day of RNAseq-defined inflammation (**Figure 6B**). Furthermore, the onset of RNAemia correlated with the timing of IgM and IgA seroconversion, but less well with the appearance of either DENV virion or NS1 specific IgG. Notably, the only immunologic parameter that inversely correlated with the magnitude of DENV RNAemia and the abundance of NS1 antigen was the abundance of NS1 opsonizing IgA antibody, although the clinical relevance of this observation is unclear. In its totality, these data suggest that the kinetics of DENV-1 elicited inflammation and immunity are largely driven by the timing of RNAemia, but that the absolute abundance of viral antigen does not necessary directly correlate with the magnitude of the subsequent DENV-specific immune response.

## DISCUSSION

In this study we characterized the clinical, immunologic and virologic features of primary DENV-1 infection in flavivirus-naïve adults. This analysis captured both the kinetics and composition of the innate, humoral, and cellular immune responses elicited by experimental DENV-1 infection, as well as the virologic and clinical features of DENV-1 infection in 9 individuals inoculated with the under-attenuated DENV-1 strain 45AZ5. The induction of both DENV virion specific humoral immunity as well as anti-NS1 specific immunity was monitored, along with the induction and antigen specificity of DENV specific cellular immunity. Furthermore, extensive RNAseq analysis was performed on whole-blood collected from study participants throughout the course of DENV-1 infection.

Several unique immunological features of primary DENV-1 infection are revealed in this analysis, most notably the robust IgA response targeting both DENV virions and DENV NS1. The presence of DENV and NS1 specific serum IgA has previously been described following both natural primary and secondary DENV infection [32, 40–42], but the kinetics and relative magnitude of these responses were previously unclear. The timing of IgA seroconversion is notable, as the fact that it appears between IgM and IgG seroconversion makes the developmental origin of the IgA-secreting B cells somewhat ambiguous. Our previous analysis of B cells circulating following natural DENV infection demonstrated that immunoglobulins expressed by DENV-elicited/IgA-expressing plasmablasts were heavily hypermutated. This suggests that these IgA class-switched cells may be derived from previously activated and hypermutated memory B cells even following primary DENV infections. The putative memory B cell origin of the DENV-reactive IgA is strengthened by the kinetics of IgA seroconversion observed in our study, as the appearance of DENV-reactive IgA only lags IgM seroconversion by a few days and significantly precedes the appearance of DENV-specific IgG.

While IgA biased immune responses have canonically been associated with mucosal pathogen infection, more recent analyses have shown a significant IgA component following infection with other non-mucosal pathogens (such as malaria) as well as following mRNA vaccination [43, 44]. This raises the possibility that a cross-reactive IgA response is an intrinsic feature of any immune response, and may draw upon the significant clonal diversity represented in the normally mucosal-restricted immune system to speed antibody secretion upon the introduction of novel non-mucosal pathogen. Furthermore, the transient nature of circulating DENV-specific IgA isotype antibodies observed in our study has several interesting epidemiological implications. Most significantly, the presence of DENV/NS1-specific IgA may be a more temporally sensitive indication of recent DENV infection than IgM or IgG alone.

Dissecting the transcriptional profile associated with acute DENV infection and identifying transcriptional signatures that correlate with clinical outcome of infection has been a longstanding goal in the dengue community [26, 27, 39]. Hanley et al provided insight into the kinetics of the inflammatory transcriptional program elicited by infection with the NIH rDEN2Δ30 strain and additionally identified pre-infection transcriptional signatures that correlated with the clinical outcomes of infection, including the development of rash following infection [27]. However, it must be noted that the DENV-1 45AZ5 DENV-1 strain used in our study generated 100-1,000-fold higher levels of viremia than the rDEN2Δ30 strain and resulted in a broader-range of dengue-like symptoms, including fever. Accordingly, the more robust clinical response to 45AZ5 infection appears to be accompanied by a quantitatively and qualitatively more vigorous inflammatory response than is observed following rDEN2Δ30 infection. When coupled with the extensive sampling performed in our analysis, the reactogenicity and immunogenicity of 45AZ5 allowed for the identification, characterization, and head-to-head comparison of the innate/inflammatory response to acute DENV infection as well as the subsequent adaptive immune response.

The DHIM tool was designed to recapitulate the human natural infection experience, and allow the early interrogation of potential countermeasures prior to advancing to field trials. Although participants do experience dengue-like symptoms, clinical lab abnormalities, viral replication, and immune responses consistent with natural infection, it is important to note a few dissimilarities. It is unclear what viral dose a mosquito delivers to a susceptible host during the feeding process. In our trial, we deliver a consistent and standardized dose which may or may not reflect what is delivered in nature. The mosquito proboscis probes a relatively superficial anatomic space in the skin as it searches for blood filled capillaries. In the DHIM, we use a needle to deliver the virus into the subcutaneous space, a method which may produce a different antigen processing pathway than what occurs in nature. Mosquitoes deliver wild type viruses, in the DHIM we use viruses which have been attenuated through cell passage or chemical exposure. Finally, in the DHIM we do not mix the virus with mosquito saliva prior to injection, which is clearly different than the natural feeding process. Future DHIM iterations may address these differences.

In conclusion, we have demonstrated the ability to safely and consistently experimentally infect people with an under-attenuated DENV strain and generated a mild dengue like illness manifesting with clinical signs and symptoms, clinical lab abnormalities, and virologic and immunologic immune responses consistent with natural DENV infections. These studies not only provide insights into previously underappreciated immune responses, but also underscore the potential value of the DHIM tool in supporting anti-dengue countermeasure development.

## MATERIALS AND METHODS

### Dengue human infection model

The Dengue Human Infection Model and associated analysis was approved by the State University of New York Upstate Medical University (SUNY-UMU) and the Department of Defense’s Human Research Protection Office. This phase 1, open-label study (ClinicalTrials.gov identifier: NCT03869060) is an expansion of a previous study evaluating a Dengue 1 Live Virus Human Infection (DENV-1-LVHC) model (ClinicalTrials.gov identifier: NCT02372175) [14]. The current study was conducted between March 2019 and February 2021 at the State University of New York, Upstate Medical University (SUNY-UMU) in Syracuse, New York. Participants received a single subcutaneous inoculation of 3.25 × 10^3^ PFU of the 45AZ5 DENV-1 infection strain virus manufactured at the WRAIR Pilot Bioproduction Facility, Silver Spring, MD (US FDA Investigational New Drug 16332). All participants were pre-screened to ensure an absence of preexisting flavivirus using the Euroimmun dengue, West Nile, and Zika IgG ELISA kits (Lübeck, Germany). The 45AZ5 DENV-1 infection strain used in this study was generated by serial passage of the parental Nauru/West Pac/1974 DENV-1 isolate through diploid fetal rhesus lung cell line (FRhL) in the presence of 5-azacytidine, followed by plaque cloning and secondary amplification in FRhL cells [30, 45]. Quantitative DENV-1 specific reverse-transcriptase polymerase chain reaction (RT-PCR) was performed using previously published techniques [46]. Serum NS1 antigen levels were quantified using a Euroimmun dengue NS1 ELISA kit according to the manufacture’s protocol (Lübeck, Germany). Day of NS1 antigenemia onset was defined as the day at which there was a detectable NS1 titer that persisted for more than one day.

### DENV-1 virion-capture ELISA

DENV-1 reactive serum IgM/IgA/IgG levels were assessed using a flavivirus capture ELISA protocol. In short, 96 well NUNC MaxSorb flat-bottom plates were coated with 2 μg/ml flavivirus group-reactive mouse monoclonal antibody 4G2 (Envigo Bioproducts, Inc.) diluted in borate saline buffer. Plates were washed and blocked with 0.25% BSA +1% Normal Goat Serum in PBS after overnight incubation. DENV-1 (strain Nauru/West Pac/1974) was captured for 2 hours, followed by extensive washing. Serum samples were serially diluted four-fold and plated in duplicate and incubated for 1 hour at room temperature on the captured virus. DENV-specific IgM/IgG/IgA levels were quantified using anti-human IgM HRP (**Seracare, 5220-0328**), anti-human IgG HRP (**Sigma-Aldrich, SAB3701362**), and anti-human IgA HRP (**Biolegend, 411,002**). Secondary antibody binding was quantified using the TMB Microwell Peroxidase Substrate System (KPL, cat. #50-76-00) and Synergy HT plate reader. DENV-1 (strains Nauru/West Pac/1974) was propagated in Vero cells and purified by ultracentrifugation through a 30% sucrose solution. End-point titers were determined as the reciprocal of the final dilution at which the optical density (OD) was greater than 3× of a control flavivirus naïve serum. Day of seroconversion was defined as the day at which a participant’s end-point titer exceeded that of their respective day 0 sample

### Anti-NS1 ELISA

To quantify DENV-1 NS1 reactive serum IgM/IgA/IgG antibodies levels, 96 well NUNC MaxSorb flat-bottom plates were coated with 2 μg/ml DENV-1 NS1 protein (Native Antigen) diluted in Carbonate/Bicarbonate buffer and incubated overnight at 4°C. Serum samples were serially diluted four-fold, plated, and incubated for 2 hours at room temperature. DENV-1 NS1 specific IgM/IgG/IgA levels were quantified using anti-human IgM HRP (**Seracare, 5220-0328**), anti-human IgG HRP (**Sigma-Aldrich, SAB3701362**), and anti-human IgA HRP (**Biolegend, 411,002**). Secondary antibody binding was quantified using the TMB Microwell Peroxidase Substrate System (KPL, cat. #50-76-00) and Synergy HT plate reader. End-point titers were determined as the reciprocal of the final dilution at which the optical density (OD) was greater than 2× of a control flavivirus naïve serum. Day of seroconversion was defined as the day at which a participant’s end-point titer exceeded that of their respective day 0 sample

### Anti-NS1 opsonization assay

DENV-1 NS1 expressing CEM.NK^R^ cells (**Supplemental Figure 2**) were stained with a 1:500 dilution of heat-inactivated serum diluted in PBS at room temperature for 30 min. Cells were extensively washed, then stained with goat anti-human IgA AF647 (2050-31, Southern Biotech), goat anti-human IgG AF467 (2040-31, Southern Biotech), or goat anti-human IgM AF647(2020-31, Southern Biotech). Flow cytometry analysis was performed on a BD LSRII instrument, and data analyzed using FlowJo v10.2 software (Treestar). Reported MFI values are background subtracted, with background defined as the signal observed staining DENV-1 NS1 expressing CEM.NK^R^ cells in the absence of serum. Day of seroconversion was defined as the day at which a participant’s background-subtracted NS1 specific MFI increased by at least 2× over their respective day 0 value.

### IFN-γ ELISPOT

Cryopreserved PBMC were thawed, washed twice, and placed in RPMI 1640 medium (Corning, Tewksbury, MA, USA) supplemented with 10% heat-inactivated fetal calf serum (Corning, 35-010-CV), L-glutamine (Lonza, Basel, Switzerland), and Penicillin/Streptomycin (Gibco, Waltham, MA, USA). Cellular viability was assessed by trypan blue exclusion and cells were resuspended at a concentration of 5 × 10^6^/mL and rested overnight at 37 °C. After resting, viable PBMC were washed, counted, and resuspended at a concentration of 1 × 10^6^/mL in complete cell culture media. Next, 100 μL of this cell suspension was mixed with 100 μL of the individual peptide pools listed in **Supplemental Table 4** and diluted to a final concentration 1 μg/mL/peptide (DMSO concentration 0.5%) in complete cell culture media. This cell and peptide mixture was loaded onto a 96-well PVDF plate coated with anti-IFN-γ (3420-2HW-Plus, Mabtech, Nacka, Sweden) and cultured overnight. Controls for each participant included 0.5% DMSO alone (negative) and anti-CD3 (positive). After overnight incubation, the ELISPOT plates were washed and stained with anti-IFN-γ-biotin followed by streptavidin-conjugated HRP (3420-2HW-Plus, Mabtech). Plates were developed using TMB substrate and read using a CTL-ImmunoSpot^®^ S6 Analyzer (Cellular Technology Limited, Shaker Heights, OH, USA). All peptide pools were tested in duplicate, and the adjusted mean was reported as the mean of the duplicate experimental wells after subtracting the mean value of the negative (DMSO only) control wells. Individuals were considered reactive to a peptide pool when the background-subtracted response was >50 spot forming cells (SFC)/10^6^ PBMC. All data were normalized based on the number of cells plated per well and are presented herein as SFC/10^6^ PBMC.

### RNA sequencing library preparation and sequencing

Whole blood was collected from all study participants using PAXgene RNA collection tubes (BD) and frozen at −20° C until analyzed. RNA was recovered from the collection tubes using the Qiagen PAXgene Blood RNA isolation Kit and sequencing libraries created using Illumina Stranded Total RNA Prep with Ribo-Zero Plus and IDT-Ilmn RNA UD Indexes SetA. Final library QC and quantification was performed using a Bioanalyizer (Agilent) and DNA 1000 reagents. Libraries were pooled at an equimolar ratio and sequenced on a 300 cycle Novaseq 6000 instrument using v1.5 S4 reagent set.

### RNA sequencing gene expression analysis

Raw reads from FASTQ files were mapped to the human reference transcriptome (Ensembl, Home sapiens, GRCh38) using Kallisto [47] version 0.46.2. Transcript-level counts and abundance data were imported and summarized in R (version 4.0.2) using the TxImport package [48] and TMM normalized using the package EdgeR [49, 50]. Differential gene expression analysis performed using linear modeling and Bayesian statistics in the R package Limma [51]. Genes with a log_2_ fold change of >2 and a Benjamini-Hochberg adjusted p-value < 0.01 were considered significant. Gene Ontology (GO) analysis was performed using gprofiler2 package [52].

### BCR clonotype identification and annotation

Raw FASTQ files were filtered to contain only pair-end reads and to remove any Illumina adaptor contamination and low-quality reads using Trimmomatic (v0.39) [53]. Pair-end reads were subsequently analyzed using MiXCR (v3.0.3) using the RNA-seq/non-targeted genomic analysis pipeline [54, 55].

### Statistical analysis

All statistical analyses other than RNAseq gene expression analysis was performed using GraphPad Prism 9 Software (GraphPad Software, La Jolla, CA). A P-value □ < □ 0.05 was considered significant.

### Data availability

The authors declare that all data supporting the findings of this study are available within this article and its Supplementary Information files, or from the corresponding author upon reasonable request. RNAseq gene expression data have been deposited in the Gene Expression Omnibus database under the accession code GSE182482.

## Acknowledgements

We gratefully acknowledge excellent technical assistance provided by Karen Gentile of the Upstate Medical University Molecular Analysis Core (MAC) and the members of the Institute for Global Health and Translational Science (IGHTS) of SUNY Upstate Medical University. We also acknowledge LTC Rick Jarman and Calli Rooney from the US Army, and the members of the dengue research team at Janssen Pharmaceutica. We also wish to thank all the study participants for making this study possible. The following reagents were obtained through BEI Resources, NIAID, NIH: Peptide Array, DENV-1 Singapore/S275/1990 E protein (NR-50710), DENV-1 Singapore/S275/1990 NS1 protein (NR-2751), DENV-1 Singapore/S275/1990 NS3 protein (NR-2752), DENV-1 Singapore/S275/1990 NS5 protein (NR-4203).

## Disclaimer

The opinions or assertions contained herein are the private views of the authors and are not to be construed as reflecting the official views of the US Army or the US Department of Defense. Material has been reviewed by the Walter Reed Army Institute of Research. There is no objection to its presentation and/or publication. The investigators have adhered to the policies for protection of human participants as prescribed in AR 70-25

## REFERENCES

1. Gubler, D.J., Aedes aegypti and Aedes aegypti-borne disease control in the 1990s: top down or bottom up. Charles Franklin Craig Lecture. Am J Trop Med Hyg, 1989. 40(6): p. 571–8.

2. Bhatt, S., et al., The global distribution and burden of dengue. Nature, 2013. 496(7446): p. 504–7.

3. Shepard, D.S., et al., Economic impact of dengue illness in the Americas. Am J Trop Med Hyg, 2011. 84(2): p. 200–7.

4. World mosquito program. Facts sheet Dengue. [cited 2021 27 April]; Available from: https://www.worldmosquitoprogram.org/sites/default/files/2020-11/WMP%20dengue_0.pdf.

5. Messina, J.P., et al., The current and future global distribution and population at risk of dengue. Nature Microbiology, 2019. 4(9): p. 1508–1515.

6. Tricou, V., et al., Safety and immunogenicity of a tetravalent dengue vaccine in children aged 2-17 years: a randomised, placebo-controlled, phase 2 trial. Lancet, 2020. 395(10234): p. 1434–1443.

7. Biswal, S., et al., Efficacy of a Tetravalent Dengue Vaccine in Healthy Children and Adolescents. N Engl J Med, 2019. 381(21): p. 2009–2019.

8. Kirkpatrick, B.D., et al., Robust and Balanced Immune Responses to All 4 Dengue Virus Serotypes Following Administration of a Single Dose of a Live Attenuated Tetravalent Dengue Vaccine to Healthy, Flavivirus-Naive Adults. J Infect Dis, 2015. 212(5): p. 702–10.

9. Kirkpatrick, B.D., et al., The live attenuated dengue vaccine TV003 elicits complete protection against dengue in a human challenge model. Sci Transl Med, 2016. 8(330): p. 330ra36.

10. Capeding, M.R., et al., Clinical efficacy and safety of a novel tetravalent dengue vaccine in healthy children in Asia: a phase 3, randomised, observer-masked, placebo-controlled trial. Lancet, 2014. 384(9951): p. 1358–65.

11. Villar, L., et al., Efficacy of a tetravalent dengue vaccine in children in Latin America. N Engl J Med, 2015. 372(2): p. 113–23.

12. World Health, O., Dengue vaccine: WHO position paper, July 2016 - recommendations. Vaccine, 2017. 35(9): p. 1200–1201.

13. Kaptein, S.J.F., et al., A pan-serotype dengue virus inhibitor targeting the NS3-NS4B interaction. Nature, 2021. 598(7881): p. 504–509.

14. Endy, T.P., et al., A Phase 1, Open-Label Assessment of a Dengue Virus-1 Live Virus Human Challenge Strain. J Infect Dis, 2021. 223(2): p. 258–267.

15. Harris, G., Dengue: a study of its mode of propagation and pathology. Med Rec New York 1902. 61: p. 204–7.

16. Harris, G., The dengue: a study of its pathology and mode of propagation. J Trop Med 1903. 6: p. 209–214.

17. Ashburnq, P. and C. Craig, Experimental investigations regarding the etiology of dengue fever. J Infect Dis, 1907. 4: p. 440–75.

18. Cleland, J.B., B. Bradley, and W. Macdonald, Further Experiments in the Etiology of Dengue Fever. J Hyg (Lond), 1919. 18(3): p. 217–54.

19. Siler, J.F., M.W. Hall, and A.P. Hitchens, Dengue: its history, epidemiology, mechanism of transmission, etiology, clinical manifestations, immunity, and prevention. Philippine Islands Bureau of Science monograph 20. Manila, Philippines: Bureau of Printing,, 1926.

20. Simmons JS, J.H. St. John, and F. Reynolds, Experimental studies of dengue. Manilla: Bureau of printing, 1931

21. Sawada, T., H. Sato, and S. Sai, On the experimental dengue infection in man. Nippon Igaku, 1943. 3324: p. 529–31.

22. Sabin, A.B. and R.W. Schlesinger, Production of Immunity to Dengue with Virus Modified by Propagation in Mice. Science, 1945. 101(2634): p. 640–2.

23. Hotta, S., Experimental studies on dengue. I. Isolation, identification and modification of the virus. J Infect Dis, 1952. 90(1): p. 1–9.

24. Sabin, A.B., Research on dengue during World War II. Am J Trop Med Hyg, 1952. 1(1): p. 30–50.

25. Yaoi, H., A summary of our studies on dengue. Yokohama Med Bull, 1958. 9(1): p. 1–20.

26. Waickman, A.T., et al., Temporally integrated single cell RNA sequencing analysis of PBMC from experimental and natural primary human DENV-1 infections. PLoS Pathog, 2021. 17(1): p. e1009240.

27. Hanley, J.P., et al., Immunotranscriptomic profiling the acute and clearance phases of a human challenge dengue virus serotype 2 infection model. Nat Commun, 2021. 12(1): p. 3054.

28. Grifoni, A., et al., Patterns of Cellular Immunity Associated with Experimental Infection with rDEN2Delta30 (Tonga/74) Support Its Suitability as a Human Dengue Virus Challenge Strain. J Virol, 2017. 91(8).

29. Repik, P.M., et al., RNA fingerprinting as a method for distinguishing dengue 1 virus strains. Am J Trop Med Hyg, 1983. 32(3): p. 577–89.

30. McKee, K.T., Jr., et al., fs. Am J Trop Med Hyg, 1987. 36(2): p. 435–42.

31. Wegman, A.D., et al., Monomeric IgA Antagonizes IgG-Mediated Enhancement of DENV Infection. Front Immunol, 2021. 12: p. 777672.

32. Waickman, A.T., et al., Transcriptional and clonal characterization of B cell plasmablast diversity following primary and secondary natural DENV infection. EBioMedicine, 2020. 54: p. 102733.

33. Winkler, G., et al., Newly synthesized dengue-2 virus nonstructural protein NS1 is a soluble protein but becomes partially hydrophobic and membrane-associated after dimerization. Virology, 1989. 171(1): p. 302–5.

34. Pryor, M.J. and P.J. Wright, The effects of site-directed mutagenesis on the dimerization and secretion of the NS1 protein specified by dengue virus. Virology, 1993. 194(2): p. 769–80.

35. Sanchez Vargas, L.A., et al., Non-structural protein 1-specific antibodies directed against Zika virus in humans mediate antibody-dependent cellular cytotoxicity. Immunology, 2021. 164(2): p. 386–397.

36. Chung, K.M., et al., Antibody recognition of cell surface-associated NS1 triggers Fc-gamma receptor-mediated phagocytosis and clearance of West Nile Virus-infected cells. J Virol, 2007. 81(17): p. 9551–5.

37. Kurane, I., et al., T-cell responses to dengue virus in humans. Trop Med Health, 2011. 39(4 Suppl): p. 45–51.

38. Waickman, A.T., et al., Assessing the Diversity and Stability of Cellular Immunity Generated in Response to the Candidate Live-Attenuated Dengue Virus Vaccine TAK-003. Front Immunol, 2019. 10: p. 1778.

39. Robinson, M., et al., A 20-Gene Set Predictive of Progression to Severe Dengue. Cell Rep, 2019. 26(5): p. 1104–1111 e4.

40. Alagarasu, K., et al., A meta-analysis of the diagnostic accuracy of dengue virus-specific IgA antibody-based tests for detection of dengue infection. Epidemiol Infect, 2016. 144(4): p. 876–86.

41. Koraka, P., et al., Kinetics of dengue virus-specific serum immunoglobulin classes and subclasses correlate with clinical outcome of infection. J Clin Microbiol, 2001. 39(12): p. 4332–8.

42. Bachal, R., et al., Higher levels of dengue-virus-specific IgG and IgA during pre-defervescence associated with primary dengue hemorrhagic fever. Arch Virol, 2015. 160(10): p. 2435–43.

43. Berry, A.A., et al., Immunoprofiles associated with controlled human malaria infection and naturally acquired immunity identify a shared IgA pre-erythrocytic immunoproteome. NPJ Vaccines, 2021. 6(1): p. 115.

44. Azzi, L., et al., Mucosal immune response in BNT162b2 COVID-19 vaccine recipients. EBioMedicine, 2021. 75: p. 103788.

45. Puri, B., et al., Molecular analysis of dengue virus attenuation after serial passage in primary dog kidney cells. J Gen Virol, 1997. 78 (Pt 9): p. 2287–91.

46. Houng, H.S., et al., Development of a fluorogenic RT-PCR system for quantitative identification of dengue virus serotypes 1-4 using conserved and serotype-specific 3’ noncoding sequences. J Virol Methods, 2001. 95(1-2): p. 19–32.

47. Bray, N.L., et al., Erratum: Near-optimal probabilistic RNA-seq quantification. Nat Biotechnol, 2016. 34(8): p. 888.

48. Soneson, C., M.I. Love, and M.D. Robinson, Differential analyses for RNA-seq: transcript-level estimates improve gene-level inferences. F1000Res, 2015. 4: p. 1521.

49. Robinson, M.D., D.J. McCarthy, and G.K. Smyth, edgeR: a Bioconductor package for differential expression analysis of digital gene expression data. Bioinformatics, 2010. 26(1): p. 139–40.

50. McCarthy, D.J., Y. Chen, and G.K. Smyth, Differential expression analysis of multifactor RNA-Seq experiments with respect to biological variation. Nucleic Acids Res, 2012. 40(10): p. 4288–97.

51. Ritchie, M.E., et al., limma powers differential expression analyses for RNA-sequencing and microarray studies. Nucleic Acids Res, 2015. 43(7): p. e47.

52. Kolberg, L., et al., gprofiler2 -- an R package for gene list functional enrichment analysis and namespace conversion toolset g:Profiler. F1000Res, 2020. 9.

53. Bolger, A.M., M. Lohse, and B. Usadel, Trimmomatic: a flexible trimmer for Illumina sequence data. Bioinformatics, 2014. 30(15): p. 2114–20.

54. Bolotin, D.A., et al., MiXCR: software for comprehensive adaptive immunity profiling. Nat Methods, 2015. 12(5): p. 380–1.

55. Bolotin, D.A., et al., Antigen receptor repertoire profiling from RNA-seq data. Nat Biotechnol, 2017. 35(10): p. 908–911.

