## Supplemental tables for "Evolution of inflammation and immunity in a dengue virus 1 human infection model"

|  | **All Participants (N=9)** |
| --- | --- |
| Age (years) |  |
| Mean (SD) | 34.6 (8.8) |
| Median | 33.0 |
| Min, Max | 20, 45 |
| Sex, n (%) |  |
| Male | 3 (33.3) |
| Female | 6 (66.7) |
| Ethnicity, n (%) |  |
| Hispanic or Latino | 3 (33.3) |
| Non-Hispanic or Latino | 6 (66.7) |
| Race, n (%) |  |
| White | 7 (77.8) |
| Black or African American | 1 (11.1) |
| American Indian or Alaska Native | 0 (0.0) |
| Asian | 0 (0.0) |
| Native Hawaiian or Other Pacific  Islander | 0 (0.0) |
| Other or Multiple [1] | 1 (11.1) |

**Supplemental Table 1. Study participants demographics**

**Supplemental Table 2. Summary of solicited Systemic Adverse Events reported within 28 days after inoculation or 7 days post hospitalization (whichever later)**

|  | Any | Mild/Moderate | Severe |
| --- | --- | --- | --- |
| Headache | 9/9 (100%) | 8/9 (88.8%) | 1/9 (11.1%) |
| Rash | 7/9 (77.7%) | 6/9 (66.6%) | 1/9 (11.1%) |
| Fever | 7/9 (77.7%) | 6/9 (66.6%) | 1/9 (11.1%) |
| Eye pain | 7/9 (77.7%) | 6/9 (66.6%) | 1/9 (11.1%) |
| Weakness/Fatigue | 6/9 (66.6%) | 3/9 (33.3%) | 3/9 (33.3%) |
| Myalgia | 7/9 (77.7%) | 7/9 (77.7%) | 0/9 (0%) |

**Supplemental Table 3. Summary of Lab Abnormalities**

| **Number of subjects experiencing at least one** |  | **Maximum intensity** | **All Subjects (N=9) n/M(%)** |
| --- | --- | --- | --- |
| Lab abnormality |  |  | 8/9(88.9) |
|  |  | Mild | 2/9(22.2) |
|  |  | Moderate | 3/9(33.3) |
|  |  | Severe | 2/9(22.2) |
|  |  | Potentially life-threatening | 1/9(11.1) |
| Blood and lymphatic system disorders |  |  | 6/9(66.7) |
|  |  | Moderate | 5/9(55.6) |
|  |  | Severe | 1/9(11.1) |
|  | Leukopenia |  | 6/9(66.7) |
|  |  | Mild | 2/9(22.2) |
|  |  | Moderate | 4/9(44.4) |
|  | Lymphopenia |  | 3/9(33.3) |
|  |  | Mild | 2/9(22.2) |
|  |  | Severe | 1/9(11.1) |
|  | Neutropenia |  | 2/9(22.2) |
|  |  | Moderate | 2/9(22.2) |
|  | Thrombocytopenia |  | 1/9(11.1) |
|  |  | Mild | 1/9(11.1) |
| Investigations |  |  | 8/9(88.9) |
|  |  | Mild | 4/9(44.4) |
|  |  | Moderate | 2/9(22.2) |
|  |  | Severe | 2/9(22.2) |
|  | Activated partial thromboplastin time |  | 1/9(11.1) |
|  |  | Mild | 1/9(11.1) |
|  | Alanine aminotransferase increased |  | 1/9(11.1) |
|  |  | Mild | 1/9(11.1) |
|  | Aspartate aminotransferase increased |  | 3/9(33.3) |
|  |  | Mild | 3/9(33.3) |
|  | Blood alkaline phosphatase increased |  | 1/9(11.1) |
|  |  | Mild | 1/9(11.1) |
|  | Blood calcium increased |  | 1/9(11.1) |
|  |  | Mild | 1/9(11.1) |
|  | Blood sodium decreased |  | 1/9(11.1) |
|  |  | Mild | 1/9(11.1) |
|  | Eosinophil count increased |  | 1/9(11.1) |
|  |  | Mild | 1/9(11.1) |
|  | Hemoglobin decreased |  | 3/9(33.3) |
|  |  | Mild | 2/9(22.2) |
|  |  | Moderate | 1/9(11.1) |
|  | Lymphocyte count decreased |  | 4/9(44.4) |
|  |  | Mild | 2/9(22.2) |
|  |  | Moderate | 2/9(22.2) |
|  | Neutrophil count decreased |  | 4/9(44.4) |
|  |  | Moderate | 2/9(22.2) |
|  |  | Severe | 2/9(22.2) |
|  | Platelet count decreased |  | 3/9(33.3) |
|  |  | Mild | 2/9(22.2) |
|  |  | Moderate | 1/9(11.1) |
|  | Protein total decreased |  | 1/9(11.1) |
|  |  | Mild | 1/9(11.1) |
|  | White blood cell count increased |  | 2/9(22.2) |
|  |  | Mild | 2/9(22.2) |
| Metabolism and nutrition disorders |  |  | 6/9(66.7) |
|  |  | Mild | 4/9(44.4) |
|  |  | Moderate | 1/9(11.1) |
|  |  | Potentially life-threatening | 1/9(11.1) |
|  | Hypocalcemia |  | 3/9(33.3) |
|  |  | Mild | 3/9(33.3) |
|  | Hypoglycemia |  | 2/9(22.2) |
|  |  | Moderate | 1/9(11.1) |
|  |  | Potentially life-threatening | 1/9(11.1) |
|  | Hyponatremia |  | 1/9(11.1) |
|  |  | Mild | 1/9(11.1) |
|  | Hypoproteinemia |  | 1/9(11.1) |
|  |  | Mild | 1/9(11.1) |

**Supplemental Table 4. Peptides used in IFN-γ ELISPOT assay**

| **Virus** | **Antigen** | **Supplier** | **Catalog #** |
| --- | --- | --- | --- |
| DENV-1: Singapore/S275/1990 | E | BEI | NR-50710 |
| DENV-1: Singapore/S275/1990 | NS1 | BEI | NR-2751 |
| DENV-1: Singapore/S275/1990 | NS3 | BEI | NR-2752 |
| DENV-1: Singapore/S275/1990 | NS5 | BEI | NR-4203 |

**Supplemental Table 5: Day 10 upregulated DEGs**

| **geneID** | **logFC** | **AveExpr** | **t** | **P.Value** | **adj.P.Val** | **B** |
| --- | --- | --- | --- | --- | --- | --- |
| SIGLEC1 | 5.28933912 | 4.9170254 | 5.57008222 | 1.45E-06 | 4.88E-04 | 5.14181002 |
| RSAD2 | 5.12713098 | 7.45523814 | 6.23112006 | 1.55E-07 | 1.86E-04 | 7.2244684 |
| USP41 | 4.97810235 | -0.3332919 | 4.14262612 | 1.54E-04 | 0.00761562 | -0.0357048 |
| IFI44L | 4.89630162 | 7.86949929 | 6.51809444 | 5.87E-08 | 1.86E-04 | 8.11704845 |
| USP18 | 4.78795711 | 4.06786798 | 4.97795746 | 1.04E-05 | 0.0015773 | 3.33039413 |
| IFIT1 | 4.7806291 | 7.96079753 | 6.14184276 | 2.10E-07 | 1.88E-04 | 6.92851822 |
| OAS3 | 4.35410419 | 8.45525895 | 6.18809519 | 1.80E-07 | 1.86E-04 | 7.06295287 |
| CMPK2 | 4.34211562 | 6.30431638 | 6.15942995 | 1.98E-07 | 1.86E-04 | 7.03550612 |
| SERPING1 | 4.33547355 | 5.10853182 | 6.01475457 | 3.23E-07 | 2.18E-04 | 6.56095313 |
| IFI6 | 4.28717795 | 7.30993194 | 6.38353384 | 9.26E-08 | 1.86E-04 | 7.72871153 |
| IFI44 | 4.24332021 | 6.48312526 | 6.17478727 | 1.88E-07 | 1.86E-04 | 7.08596531 |
| IFIT3 | 4.24240159 | 8.51353137 | 6.18170763 | 1.83E-07 | 1.86E-04 | 7.03131762 |
| OASL | 4.13383203 | 6.65230664 | 6.06557972 | 2.72E-07 | 2.18E-04 | 6.74205204 |
| EXOC3L1 | 4.08517533 | 1.57737062 | 4.32232158 | 8.71E-05 | 0.00563067 | 1.27047967 |
| IFIT2 | 3.95768905 | 8.9735811 | 5.7238075 | 8.62E-07 | 3.84E-04 | 5.58321496 |
| IFITM3 | 3.94580746 | 8.11792354 | 6.02510258 | 3.12E-07 | 2.18E-04 | 6.58410934 |
| BATF2 | 3.93186641 | 4.5098331 | 5.21608921 | 4.73E-06 | 9.00E-04 | 4.07268239 |
| HERC5 | 3.92934358 | 6.79190049 | 5.85009111 | 5.63E-07 | 3.02E-04 | 6.0617348 |
| ATF3 | 3.91736562 | 2.14500789 | 4.19364889 | 1.31E-04 | 0.00705385 | 0.98340754 |
| LAMP3 | 3.82469431 | 3.64792721 | 4.49226025 | 5.06E-05 | 0.0041574 | 1.88794784 |
| ETV7 | 3.77868209 | 3.95378733 | 4.68829819 | 2.69E-05 | 0.00280203 | 2.46821467 |
| OAS1 | 3.66964493 | 7.42875753 | 6.19344668 | 1.76E-07 | 1.86E-04 | 7.14092201 |
| SPATS2L | 3.60105974 | 5.24305999 | 5.20918701 | 4.83E-06 | 9.00E-04 | 4.05441889 |
| MT2A | 3.58791401 | 5.30056281 | 5.48705511 | 1.91E-06 | 5.53E-04 | 4.92239718 |
| MX1 | 3.54868417 | 8.62072361 | 6.18756549 | 1.80E-07 | 1.86E-04 | 7.05207278 |
| RP11-466G12.3 | 3.52138707 | -0.420473 | 4.28550866 | 9.79E-05 | 0.00603748 | 0.65618487 |
| RP4-641G12.3 | 3.49335065 | 0.00407114 | 4.71167228 | 2.49E-05 | 0.00270876 | 1.79151158 |
| EPSTI1 | 3.40483944 | 6.79024856 | 6.86913068 | 1.79E-08 | 1.86E-04 | 9.29866305 |
| CARD17 | 3.38974587 | -0.4710274 | 4.69348239 | 2.64E-05 | 0.00277627 | 1.33701578 |
| ZDHHC4P1 | 3.29123309 | 3.19294297 | 5.85414972 | 5.55E-07 | 3.02E-04 | 5.95634128 |
| LY6E | 3.29096159 | 8.27192948 | 6.04107531 | 2.95E-07 | 2.18E-04 | 6.64022358 |
| FBXO39 | 3.11623692 | 1.19699362 | 5.48977715 | 1.89E-06 | 5.53E-04 | 4.27831017 |
| OAS2 | 3.08162882 | 8.04770319 | 6.00227119 | 3.37E-07 | 2.18E-04 | 6.52288643 |
| SMTNL1 | 2.90961277 | 4.80724124 | 5.32190721 | 3.32E-06 | 7.35E-04 | 4.40540173 |
| RMI2 | 2.87018557 | 2.72868882 | 4.22400741 | 1.19E-04 | 0.00670342 | 1.11489591 |
| IL1RN | 2.86942241 | 7.29093357 | 4.66917458 | 2.86E-05 | 0.00288994 | 2.40043803 |
| EIF2AK2 | 2.86661196 | 8.23836112 | 6.22919419 | 1.56E-07 | 1.86E-04 | 7.21898947 |
| RTP4 | 2.86068258 | 4.69971644 | 5.41031669 | 2.47E-06 | 6.28E-04 | 4.68217208 |
| LAP3 | 2.8578458 | 6.47855507 | 6.31916147 | 1.15E-07 | 1.86E-04 | 7.56553814 |
| DHX58 | 2.84179357 | 5.59355218 | 5.37611892 | 2.77E-06 | 6.82E-04 | 4.57295317 |
| IFI35 | 2.79696682 | 5.86924027 | 5.79975403 | 6.67E-07 | 3.30E-04 | 5.91350511 |
| IFIH1 | 2.79295036 | 4.64938461 | 5.36988461 | 2.83E-06 | 6.82E-04 | 4.55541731 |
| PLSCR2 | 2.79206355 | 2.74175719 | 5.44173739 | 2.22E-06 | 6.00E-04 | 4.71050799 |
| PLSCR1 | 2.78667279 | 7.14933055 | 5.86936561 | 5.27E-07 | 3.02E-04 | 6.12773211 |
| DDX60 | 2.77741734 | 6.84319897 | 6.69896506 | 3.18E-08 | 1.86E-04 | 8.76843931 |
| TMPRSS2 | 2.76277301 | -1.6361692 | 3.96963617 | 2.63E-04 | 0.0099939 | -0.7484047 |
| C1QB | 2.75879546 | 1.04818111 | 4.5326293 | 4.45E-05 | 0.00383522 | 1.66260605 |
| IRF7 | 2.74916201 | 7.13751945 | 5.70257159 | 9.26E-07 | 3.87E-04 | 5.6019334 |
| DDX58 | 2.72662342 | 8.17224372 | 5.7090694 | 9.06E-07 | 3.87E-04 | 5.5980808 |
| TNFAIP6 | 2.7110423 | 5.01580247 | 4.22364232 | 1.19E-04 | 0.00670342 | 1.05002071 |
| GBP1 | 2.67649798 | 7.7556123 | 6.25580794 | 1.43E-07 | 1.86E-04 | 7.33849815 |
| SAMD9L | 2.67405473 | 8.53431501 | 6.19817362 | 1.73E-07 | 1.86E-04 | 7.10325287 |
| ANKRD22 | 2.6621509 | 3.81398993 | 4.71365797 | 2.47E-05 | 0.00270876 | 2.54367511 |
| IFIT5 | 2.65846346 | 6.76290143 | 5.73934207 | 8.18E-07 | 3.84E-04 | 5.72028608 |
| AC007919.18 | 2.59468385 | 1.11313906 | 4.34630963 | 8.07E-05 | 0.00542228 | 1.32104624 |
| UBE2L6 | 2.57609111 | 7.70048281 | 6.24474398 | 1.48E-07 | 1.86E-04 | 7.3017434 |
| LIPA | 2.57040806 | 6.30864899 | 6.17654797 | 1.87E-07 | 1.86E-04 | 7.11402157 |
| PNPT1 | 2.56839029 | 5.45939854 | 5.4402196 | 2.24E-06 | 6.00E-04 | 4.77387191 |
| RUFY4 | 2.56798385 | 2.08182909 | 4.1809425 | 1.36E-04 | 0.00708214 | 0.96546791 |
| OR52M2P | 2.56266818 | -0.6309252 | 4.4075146 | 6.64E-05 | 0.00484142 | 0.78131702 |
| LLpac-136A2.1 | 2.558184 | -0.9240924 | 4.49301703 | 5.05E-05 | 0.0041574 | 1.06079828 |
| AC007899.3 | 2.55502032 | 1.8636924 | 4.76276373 | 2.11E-05 | 0.0024314 | 2.60263196 |
| TREX1 | 2.52256094 | 4.97323551 | 5.27924989 | 3.83E-06 | 7.92E-04 | 4.26900792 |
| XAF1 | 2.51056562 | 8.38015327 | 6.48873778 | 6.49E-08 | 1.86E-04 | 8.02805586 |
| TNFSF13B | 2.51044741 | 7.09149607 | 5.07681897 | 7.50E-06 | 0.00122672 | 3.64248391 |
| PARP14 | 2.49209437 | 9.57426596 | 6.00365384 | 3.35E-07 | 2.18E-04 | 6.4252801 |
| TRIM6 | 2.48970255 | 2.6111843 | 5.42851012 | 2.32E-06 | 6.07E-04 | 4.66093925 |
| AC003080.4 | 2.48752958 | 0.5368816 | 4.17916776 | 1.37E-04 | 0.00708214 | 0.76640666 |
| FRMD3 | 2.48321788 | 4.33959254 | 4.62282128 | 3.32E-05 | 0.00318747 | 2.26094863 |
| C1QC | 2.45746823 | 0.24040917 | 4.16529332 | 1.43E-04 | 0.00726978 | 0.54163624 |
| ZBP1 | 2.44186533 | 7.24901157 | 5.55903188 | 1.50E-06 | 4.88E-04 | 5.14871718 |
| HERC6 | 2.43853919 | 6.1327843 | 5.86156199 | 5.42E-07 | 3.02E-04 | 6.11083487 |
| SCO2 | 2.43733028 | 3.41575559 | 5.2131069 | 4.77E-06 | 9.00E-04 | 4.05485603 |
| CD274 | 2.39357619 | 5.11770943 | 4.39045207 | 7.02E-05 | 0.00501851 | 1.53686814 |
| RP11-820K3.2 | 2.3832536 | 1.44697259 | 4.48520469 | 5.18E-05 | 0.00421344 | 1.77202663 |
| GBP4 | 2.36426952 | 7.58162532 | 5.55794533 | 1.51E-06 | 4.88E-04 | 5.14464536 |
| TNFSF10 | 2.34513832 | 8.55380545 | 5.13142533 | 6.26E-06 | 0.00110015 | 3.80580503 |
| PARP9 | 2.32844771 | 8.40541098 | 6.28634359 | 1.29E-07 | 1.86E-04 | 7.39561422 |
| ACO1 | 2.31954731 | 7.13833425 | 5.45890697 | 2.10E-06 | 5.81E-04 | 4.8346327 |
| TFEC | 2.29603777 | 5.89552158 | 4.38625234 | 7.11E-05 | 0.00504463 | 1.51680605 |
| TCN2 | 2.29510324 | 2.79259981 | 4.07342925 | 1.91E-04 | 0.00845425 | 0.69172565 |
| SAMD4A | 2.28626494 | 4.46504118 | 4.85439448 | 1.56E-05 | 0.00201151 | 2.95694848 |
| RNA5SP39 | 2.28149067 | 1.07322946 | 4.01598103 | 2.28E-04 | 0.00925247 | 0.46867615 |
| HELZ2 | 2.27785449 | 7.78274997 | 5.15008094 | 5.88E-06 | 0.00104384 | 3.87181563 |
| AC074338.4 | 2.27066556 | 2.09771067 | 4.86550906 | 1.51E-05 | 0.00197997 | 2.94902764 |
| RPL37P6 | 2.2608075 | 0.43198787 | 4.84870165 | 1.59E-05 | 0.0020215 | 2.36584541 |
| STAT2 | 2.23860855 | 7.65709432 | 5.59106506 | 1.35E-06 | 4.78E-04 | 5.24702284 |
| BST2 | 2.23841456 | 5.994799 | 5.64578523 | 1.12E-06 | 4.05E-04 | 5.4240373 |
| STAT1 | 2.22743235 | 8.03222344 | 6.81241515 | 2.17E-08 | 1.86E-04 | 9.07584338 |
| FBXO6 | 2.21469778 | 4.61604365 | 5.54829488 | 1.56E-06 | 4.96E-04 | 5.11614319 |
| PML | 2.20323024 | 7.91716837 | 5.64770958 | 1.11E-06 | 4.05E-04 | 5.41802096 |
| C1QA | 2.17987978 | 1.94303161 | 5.17782772 | 5.37E-06 | 9.80E-04 | 3.75029915 |
| TDRD7 | 2.16817935 | 5.96333466 | 5.72681862 | 8.53E-07 | 3.84E-04 | 5.68172493 |
| DDX60L | 2.1641171 | 8.70357217 | 5.51104539 | 1.76E-06 | 5.43E-04 | 4.96787376 |
| TTC26 | 2.13263162 | 3.36703253 | 4.5043839 | 4.87E-05 | 0.00405157 | 1.91924751 |
| PARP12 | 2.12281677 | 7.23577109 | 5.6699239 | 1.03E-06 | 4.05E-04 | 5.49908191 |
| NUDT19P5 | 2.11773239 | -0.4214855 | 4.87649394 | 1.45E-05 | 0.00197659 | 1.97291785 |
| AIM2 | 2.10535452 | 4.53893729 | 5.3577762 | 2.95E-06 | 6.89E-04 | 4.51681509 |
| MTND4P26 | 2.09894625 | 1.56855523 | 4.41214469 | 6.55E-05 | 0.00479797 | 1.60028611 |
| TOR1B | 2.0858713 | 6.17430835 | 5.16725915 | 5.56E-06 | 9.95E-04 | 3.91357441 |
| TTC21A | 2.07430352 | 4.29185633 | 4.36766321 | 7.54E-05 | 0.00515801 | 1.48063394 |
| ZCCHC2 | 2.072024 | 7.79786636 | 5.7182061 | 8.78E-07 | 3.84E-04 | 5.64252896 |
| KIAA1958 | 2.06820903 | 4.8600693 | 4.82518502 | 1.72E-05 | 0.00211912 | 2.85497093 |
| TRIM22 | 2.06430117 | 9.02455307 | 5.80842956 | 6.48E-07 | 3.29E-04 | 5.87180208 |
| MS4A4A | 2.06027982 | 2.69171556 | 4.85436158 | 1.56E-05 | 0.00201151 | 2.95676503 |
| EPHB2 | 2.0596664 | 2.89674599 | 5.06652006 | 7.76E-06 | 0.00124757 | 3.59107029 |
| SAMD9 | 2.03594466 | 8.32363289 | 4.31590608 | 8.89E-05 | 0.00566862 | 1.36751633 |
| LGALS9 | 2.03326241 | 7.61471601 | 5.28604893 | 3.74E-06 | 7.92E-04 | 4.29358823 |
| CCR1 | 2.03289549 | 7.57595165 | 4.71383975 | 2.47E-05 | 0.00270876 | 2.5342753 |
| XXbac-BPG541D20.6 | 2.0174689 | -0.1495043 | 4.76601447 | 2.09E-05 | 0.00242058 | 2.00609701 |

**Supplemental Table 6: Day 10 downregulated DEGs**

| **geneID** | **logFC** | **AveExpr** | **t** | **P.Value** | **adj.P.Val** | **B** |
| --- | --- | --- | --- | --- | --- | --- |
| ADAMTS5 | -2.5713877 | -0.0678239 | -4.2878975 | 9.72E-05 | 0.00603152 | 0.86656164 |
| CXCL6 | -2.0613406 | -0.6719037 | -4.2141531 | 1.23E-04 | 0.00676507 | 0.5176462 |

**Supplemental Table 7: Day 14 upregulated DEGs**

| **geneID** | **logFC** | **AveExpr** | **t** | **P.Value** | **adj.P.Val** | **B** |
| --- | --- | --- | --- | --- | --- | --- |
| IFI27 | 7.93347021 | 4.73260463 | 6.341738 | 1.07E-07 | 4.27E-05 | 7.5798908 |
| SIGLEC1 | 5.45400918 | 4.9170254 | 5.69134883 | 9.62E-07 | 1.56E-04 | 5.56889057 |
| IFI44L | 4.80809266 | 7.86949929 | 6.40070019 | 8.74E-08 | 4.01E-05 | 7.83929441 |
| CXCL11 | 4.74554915 | -1.799615 | 3.90327068 | 3.22E-04 | 0.00681958 | -0.6997596 |
| USP18 | 4.340308 | 4.06786798 | 4.52697666 | 4.53E-05 | 0.00175724 | 1.95715788 |
| C1QC | 4.21337769 | 0.24040917 | 7.62333764 | 1.42E-09 | 4.46E-06 | 10.3192292 |
| SERPING1 | 4.2011929 | 5.10853182 | 5.82846665 | 6.06E-07 | 1.26E-04 | 6.00329553 |
| RSAD2 | 4.14003508 | 7.45523814 | 5.3640527 | 2.88E-06 | 2.97E-04 | 4.51172107 |
| PXT1 | 3.92190445 | -0.6017883 | 3.95738208 | 2.73E-04 | 0.00606092 | -0.0619256 |
| C1QB | 3.85243664 | 1.04818111 | 6.60362981 | 4.40E-08 | 2.67E-05 | 7.86615664 |
| IFI44 | 3.83893165 | 6.48312526 | 5.69018305 | 9.65E-07 | 1.56E-04 | 5.5468332 |
| NEURL3 | 3.71155614 | -0.7379014 | 3.89810266 | 3.28E-04 | 0.00689636 | -0.1366333 |
| IFITM3 | 3.66232202 | 8.11792354 | 5.6663928 | 1.05E-06 | 1.65E-04 | 5.48526987 |
| ATF3 | 3.64796173 | 2.14500789 | 3.89374333 | 3.32E-04 | 0.0069656 | 0.19599576 |
| GBP1P1 | 3.62281344 | 2.29778179 | 5.03639548 | 8.58E-06 | 5.83E-04 | 3.51517375 |
| KCTD14 | 3.62273191 | -0.4284207 | 5.70427733 | 9.21E-07 | 1.56E-04 | 4.82134665 |
| BATF2 | 3.59416766 | 4.5098331 | 4.77754478 | 2.01E-05 | 0.00106378 | 2.68675147 |
| CDC25A | 3.58666266 | -0.4712332 | 4.48575326 | 5.17E-05 | 0.00193272 | 1.54067155 |
| OAS3 | 3.58002207 | 8.45525895 | 5.35310664 | 2.99E-06 | 2.99E-04 | 4.49031733 |
| CMPK2 | 3.54113036 | 6.30431638 | 5.23262184 | 4.47E-06 | 3.75E-04 | 4.0769934 |
| LY6E | 3.4745293 | 8.27192948 | 6.25290852 | 1.44E-07 | 5.21E-05 | 7.36538552 |
| IFI6 | 3.42899215 | 7.30993194 | 5.38383309 | 2.70E-06 | 2.84E-04 | 4.56402743 |
| FBXO39 | 3.42763138 | 1.19699362 | 6.14165384 | 2.10E-07 | 6.81E-05 | 6.57075929 |
| ETV7 | 3.42189499 | 3.95378733 | 4.23683334 | 1.14E-04 | 0.00336633 | 1.07258835 |
| CARD17 | 3.41525738 | -0.4710274 | 4.76773145 | 2.07E-05 | 0.00108017 | 2.0629111 |
| SPATS2L | 3.40515225 | 5.24305999 | 4.93723177 | 1.19E-05 | 7.36E-04 | 3.15101832 |
| OAS1 | 3.37912056 | 7.42875753 | 5.78209379 | 7.08E-07 | 1.34E-04 | 5.84596637 |
| EPSTI1 | 3.34691563 | 6.79024856 | 6.73052023 | 2.86E-08 | 1.92E-05 | 8.91210078 |
| HNRNPCP4 | 3.34193976 | -2.3263868 | 4.13737686 | 1.56E-04 | 0.0041641 | -0.1049377 |
| IFIT1 | 3.29759912 | 7.96079753 | 4.68855623 | 2.68E-05 | 0.00126813 | 2.38961616 |
| PBK | 3.28709593 | -1.3878625 | 5.15199892 | 5.85E-06 | 4.64E-04 | 2.9308033 |
| MT2A | 3.28352516 | 5.30056281 | 5.05332235 | 8.11E-06 | 5.67E-04 | 3.51374689 |
| RMI2 | 3.27812218 | 2.72868882 | 4.91848449 | 1.27E-05 | 7.70E-04 | 3.16007102 |
| LIF | 3.22151501 | -0.376349 | 4.33924292 | 8.26E-05 | 0.00268178 | 1.19308314 |
| ANKRD22 | 3.20890316 | 3.81398993 | 5.73658967 | 8.26E-07 | 1.44E-04 | 5.7114729 |
| IFIT3 | 3.20749187 | 8.51353137 | 5.00316486 | 9.57E-06 | 6.38E-04 | 3.3895502 |
| HIST1H3G | 3.20584982 | 3.29216642 | 6.09760875 | 2.44E-07 | 7.06E-05 | 6.8670601 |
| HIST1H3C | 3.20386106 | 2.45151343 | 5.42169592 | 2.38E-06 | 2.69E-04 | 4.71736987 |
| RP4-641G12.3 | 3.17638213 | 0.00407114 | 4.26339543 | 1.05E-04 | 0.00318019 | 1.01365267 |
| OASL | 3.17530057 | 6.65230664 | 4.91288051 | 1.29E-05 | 7.74E-04 | 3.05844898 |
| ZDHHC4P1 | 3.1599642 | 3.19294297 | 5.62013265 | 1.22E-06 | 1.81E-04 | 5.33676624 |
| TMPRSS2 | 3.15885408 | -1.6361692 | 4.68929343 | 2.68E-05 | 0.00126813 | 1.36984309 |
| IGLV3-25 | 3.15557291 | 2.39379255 | 4.21099401 | 1.24E-04 | 0.00356262 | 1.03359306 |
| IGLV3-1 | 3.14158154 | 3.31382776 | 4.62512564 | 3.30E-05 | 0.00143541 | 2.23506471 |
| HIST1H2BM | 3.0774712 | 1.84300985 | 5.54985098 | 1.55E-06 | 2.06E-04 | 5.0927266 |
| CXCL9 | 3.02641038 | -1.0610622 | 5.10144991 | 6.91E-06 | 5.22E-04 | 2.8215599 |
| RUFY4 | 3.01924409 | 2.08182909 | 5.04843908 | 8.24E-06 | 5.72E-04 | 3.54287483 |
| RRM2 | 2.99987922 | 2.90016402 | 5.27917218 | 3.83E-06 | 3.43E-04 | 4.25903124 |
| NAT8 | 2.99563832 | 0.03652986 | 3.8592481 | 3.69E-04 | 0.0074414 | 0.04162973 |
| HIST1H2AB | 2.97167184 | 1.7752598 | 5.29866894 | 3.59E-06 | 3.32E-04 | 4.31915737 |
| HIST1H2BO | 2.95486391 | 3.25612525 | 7.460064 | 2.46E-09 | 4.62E-06 | 11.2114506 |
| ANO5 | 2.9519081 | -0.2939582 | 3.86882214 | 3.58E-04 | 0.00731429 | -0.0676179 |
| C1QA | 2.92950279 | 1.94303161 | 7.25242186 | 4.93E-09 | 5.45E-06 | 10.263952 |
| CDCA5 | 2.92737311 | 0.37438001 | 4.90432507 | 1.33E-05 | 7.86E-04 | 2.98669434 |
| HIST1H3J | 2.88517315 | 2.75115951 | 5.75240297 | 7.83E-07 | 1.44E-04 | 5.76319456 |
| CDK1 | 2.85213247 | 0.60997238 | 4.98806735 | 1.01E-05 | 6.59E-04 | 3.25061427 |
| NMRAL1P1 | 2.84927361 | -1.0573799 | 5.25042851 | 4.21E-06 | 3.62E-04 | 3.41969183 |
| SDC1 | 2.84626983 | -1.9419671 | 3.77094498 | 4.82E-04 | 0.00883699 | -0.5075626 |
| E2F8 | 2.81706254 | 0.67739709 | 6.13297071 | 2.16E-07 | 6.89E-05 | 6.58332283 |
| GBP1 | 2.80142465 | 7.7556123 | 6.45256126 | 7.33E-08 | 3.53E-05 | 8.01178897 |
| IGLV3-21 | 2.79145015 | 3.10743434 | 4.16458389 | 1.43E-04 | 0.00392745 | 0.86439568 |
| SMTNL1 | 2.78945799 | 4.80724124 | 5.09071078 | 7.17E-06 | 5.28E-04 | 3.62952385 |
| OAS2 | 2.78100523 | 8.04770319 | 5.49332293 | 1.87E-06 | 0.00023297 | 4.92888067 |
| RTP4 | 2.75688239 | 4.69971644 | 5.19953661 | 4.99E-06 | 4.10E-04 | 3.97563802 |
| HMMR | 2.75514391 | 0.38837942 | 5.2102837 | 4.82E-06 | 3.99E-04 | 3.86853945 |
| HIST1H2AH | 2.74593669 | 2.84712462 | 5.79223464 | 6.84E-07 | 1.31E-04 | 5.89011409 |
| ELOVL3 | 2.74532657 | -1.4171679 | 4.93821633 | 1.19E-05 | 7.36E-04 | 2.0831104 |
| HIST1H3B | 2.73439797 | 3.67190942 | 5.37937864 | 2.74E-06 | 2.86E-04 | 4.56053934 |
| CDCA2 | 2.70995561 | 0.17140536 | 4.56272002 | 4.04E-05 | 0.00162129 | 2.01550843 |
| IGLV6-57 | 2.70544753 | 1.24851897 | 5.09305269 | 7.11E-06 | 5.28E-04 | 3.65360665 |
| HIST1H2AJ | 2.68895199 | 3.13381681 | 5.00323027 | 9.57E-06 | 6.38E-04 | 3.39062359 |
| HIST1H3F | 2.66728673 | 2.9713132 | 5.27798958 | 3.84E-06 | 3.43E-04 | 4.25848239 |
| CDC45 | 2.65694375 | -0.5515261 | 3.93330133 | 2.94E-04 | 0.00635432 | 0.19530622 |
| IGLC2 | 2.65576702 | 6.06964726 | 5.80951568 | 6.46E-07 | 1.29E-04 | 5.9195259 |
| KIFC1 | 2.6512259 | 0.753573 | 4.56960938 | 3.95E-05 | 0.00159929 | 2.09018593 |
| TCN2 | 2.6359242 | 2.79259981 | 4.77360945 | 2.03E-05 | 0.00107151 | 2.71327244 |
| HIST1H2AL | 2.63478516 | 2.87666973 | 5.80127195 | 6.64E-07 | 1.31E-04 | 5.91887865 |
| DDX60 | 2.63160372 | 6.84319897 | 6.36235864 | 9.95E-08 | 4.16E-05 | 7.71588965 |
| CEP55 | 2.62465002 | 0.24269623 | 4.49122592 | 5.08E-05 | 0.00191043 | 1.79725041 |
| CENPA | 2.62220982 | -0.5000022 | 4.67130538 | 2.84E-05 | 0.0013116 | 2.05971841 |
| LAP3 | 2.60636355 | 6.47855507 | 5.81354883 | 6.37E-07 | 1.29E-04 | 5.92928426 |
| CCNB2 | 2.55281919 | 1.153791 | 5.17281437 | 5.46E-06 | 4.40E-04 | 3.87965304 |
| ANXA10 | 2.55114597 | -0.7950981 | 4.47423833 | 5.36E-05 | 0.00197026 | 1.3901627 |
| IGHV5-51 | 2.52056322 | 2.23118049 | 3.98637889 | 2.50E-04 | 0.00566285 | 0.39277191 |
| RPL37P6 | 2.50716098 | 0.43198787 | 5.48221105 | 1.94E-06 | 2.39E-04 | 4.4984345 |
| SOCS1 | 2.48969411 | 3.20970579 | 3.73927734 | 5.31E-04 | 0.00925892 | -0.3305986 |
| HIST1H1B | 2.48460985 | 4.67577083 | 5.08454689 | 7.31E-06 | 5.35E-04 | 3.59069491 |
| IGLV8-61 | 2.48393278 | 0.92428013 | 3.96343543 | 2.68E-04 | 0.00600473 | 0.38820999 |
| KIF4A | 2.48250373 | 0.42739737 | 5.91421353 | 4.53E-07 | 1.05E-04 | 5.895672 |
| MKI67 | 2.48119964 | 4.16898361 | 6.11135876 | 2.33E-07 | 7.06E-05 | 6.9054296 |
| MX1 | 2.46365507 | 8.62072361 | 4.60071816 | 3.57E-05 | 0.00151829 | 2.14568607 |
| SF3A3P2 | 2.44946737 | -0.8549619 | 4.05696115 | 2.01E-04 | 0.00488035 | 0.24742754 |
| HERC5 | 2.44786335 | 6.79190049 | 3.97596417 | 2.58E-04 | 0.00580581 | 0.19615984 |
| FRMD3 | 2.43988735 | 4.33959254 | 4.53581959 | 4.40E-05 | 0.00172402 | 1.92180146 |
| OR52M2P | 2.42230052 | -0.6309252 | 4.17685876 | 1.38E-04 | 0.00381462 | 0.64592261 |
| IGLC1 | 2.4169643 | 4.89021051 | 4.66546751 | 2.89E-05 | 0.0013211 | 2.27118056 |
| HIST2H3A | 2.40927908 | 4.0064933 | 7.49847749 | 2.16E-09 | 4.51E-06 | 11.3801073 |
| HIST2H3C | 2.40927908 | 4.0064933 | 7.49847749 | 2.16E-09 | 4.51E-06 | 11.3801073 |
| SCO2 | 2.4031521 | 3.41575559 | 5.15555476 | 5.78E-06 | 4.64E-04 | 3.87660083 |
| PYCR1 | 2.39699261 | -0.2487181 | 4.68348782 | 2.73E-05 | 0.00127843 | 2.1567944 |
| TOP2A | 2.39446444 | 2.21144413 | 5.05498626 | 8.06E-06 | 5.66E-04 | 3.5785904 |
| MYBL2 | 2.38264305 | 3.54423285 | 6.40091402 | 8.73E-08 | 4.01E-05 | 7.84408517 |
| IGKV2-28 | 2.37656221 | 2.10782537 | 4.06689602 | 1.95E-04 | 0.00480021 | 0.64360766 |
| IGLV1-40 | 2.37181384 | 3.24791198 | 4.99299494 | 9.90E-06 | 6.53E-04 | 3.3604009 |
| CXCR2P1 | 2.37114701 | 5.76999796 | 4.98088688 | 1.03E-05 | 6.73E-04 | 3.25297916 |
| ASPM | 2.36322039 | 2.54990206 | 5.97900808 | 3.64E-07 | 8.95E-05 | 6.4717111 |
| IFIT2 | 2.36135241 | 8.9735811 | 3.79221646 | 4.52E-04 | 0.00850171 | -0.2202673 |
| DTL | 2.35459427 | 1.77509196 | 5.45404192 | 2.13E-06 | 2.53E-04 | 4.79772738 |
| IRF7 | 2.33647346 | 7.13751945 | 4.95194308 | 1.13E-05 | 7.17E-04 | 3.18148757 |
| GBP4 | 2.32928409 | 7.58162532 | 5.44733441 | 2.18E-06 | 2.57E-04 | 4.77023735 |
| HIST1H2BE | 2.32485743 | 4.90197601 | 6.99492442 | 1.17E-08 | 1.00E-05 | 9.76920883 |
| GBP5 | 2.29405186 | 8.80186475 | 5.75493002 | 7.76E-07 | 1.44E-04 | 5.77123863 |
| HIST1H2BL | 2.29154495 | 2.34535236 | 4.91319103 | 1.29E-05 | 7.74E-04 | 3.14018179 |
| IGHV2-5 | 2.29138253 | 1.35632262 | 3.7513746 | 5.12E-04 | 0.00911483 | -0.2018775 |
| PDCD1LG2 | 2.27618761 | 1.33705307 | 5.46704388 | 2.04E-06 | 2.46E-04 | 4.77098672 |
| BIRC5 | 2.26637619 | 1.33988256 | 5.1791195 | 5.34E-06 | 4.33E-04 | 3.92864161 |
| RNASE1 | 2.26322099 | -1.1156337 | 4.06399517 | 1.96E-04 | 0.0048164 | 0.28183219 |
| IFI35 | 2.2533589 | 5.86924027 | 4.76502973 | 2.09E-05 | 0.00108277 | 2.57336139 |
| NUSAP1 | 2.24772526 | 1.53053638 | 5.93417832 | 4.24E-07 | 9.96E-05 | 6.26675065 |
| E2F1 | 2.23690526 | 2.14998264 | 6.48292796 | 6.61E-08 | 3.36E-05 | 8.04682839 |
| TPX2 | 2.23638835 | 2.37656719 | 6.50932828 | 6.05E-08 | 3.25E-05 | 8.14986064 |
| IGLV2-23 | 2.23109637 | 2.42257639 | 4.53240663 | 4.45E-05 | 0.00173585 | 1.9882958 |
| IGLV2-8 | 2.22419485 | 2.72432305 | 4.30350599 | 9.25E-05 | 0.00290232 | 1.2753145 |
| JCHAIN | 2.21378639 | 6.67502835 | 4.56451743 | 4.01E-05 | 0.00161536 | 1.9884861 |
| IGLV1-47 | 2.19944459 | 2.85653458 | 4.67781737 | 2.78E-05 | 0.00129048 | 2.4030072 |
| XAF1 | 2.19179007 | 8.38015327 | 5.75195803 | 7.84E-07 | 1.44E-04 | 5.75923638 |
| UBE2L6 | 2.18142464 | 7.70048281 | 5.39220281 | 2.63E-06 | 2.82E-04 | 4.59484 |
| MCM10 | 2.17980912 | 1.90780566 | 5.65080497 | 1.10E-06 | 1.68E-04 | 5.42407355 |
| MZB1 | 2.16448141 | 3.88358825 | 5.04184142 | 8.42E-06 | 5.78E-04 | 3.47682208 |
| FBXO6 | 2.15631868 | 4.61604365 | 5.38798528 | 2.66E-06 | 2.83E-04 | 4.56970551 |
| HIST1H2BI | 2.15527456 | 3.40077437 | 5.25744822 | 4.12E-06 | 3.57E-04 | 4.17632757 |
| IGLV1-51 | 2.15238909 | 3.04547045 | 3.89832205 | 3.27E-04 | 0.00689636 | 0.08069319 |
| FAM72B | 2.15158872 | 0.38083027 | 4.71775433 | 2.44E-05 | 0.00119523 | 2.38522852 |
| IGKV1D-16 | 2.14357533 | -1.5853255 | 4.16033336 | 1.45E-04 | 0.0039594 | 0.50064004 |
| TYMS | 2.13716394 | 2.20858967 | 5.30215444 | 3.55E-06 | 3.30E-04 | 4.34331106 |
| AURKB | 2.12675021 | 1.51096515 | 4.20416008 | 1.27E-04 | 0.00360699 | 1.05964053 |
| PLSCR1 | 2.09710507 | 7.14933055 | 4.58688566 | 3.73E-05 | 0.00156202 | 2.04201501 |
| EIF2AK2 | 2.09095881 | 8.23836112 | 4.75386743 | 2.17E-05 | 0.00109972 | 2.60240754 |
| ANLN | 2.08714183 | 1.96538302 | 7.00937931 | 1.12E-08 | 1.00E-05 | 9.5900875 |
| STAT1 | 2.0851279 | 8.03222344 | 6.3906954 | 9.04E-08 | 4.05E-05 | 7.81200281 |
| IGLV1-44 | 2.06923126 | 2.87436205 | 4.85862158 | 1.54E-05 | 8.70E-04 | 2.95966966 |
| MT1E | 2.06090682 | 0.86445192 | 4.72701158 | 2.37E-05 | 0.00116592 | 2.54088305 |
| BUB1 | 2.05225794 | 2.31935214 | 8.6093627 | 5.54E-11 | 5.21E-07 | 14.6366238 |
| NCAPG | 2.03992264 | 2.20751672 | 6.36554892 | 9.84E-08 | 4.16E-05 | 7.69128653 |
| IGHV4-39 | 2.02443772 | 1.79957613 | 4.36429848 | 7.63E-05 | 0.00253311 | 1.50861276 |
| TMEM51 | 2.01862189 | 0.50255538 | 4.39776028 | 6.85E-05 | 0.00235149 | 1.53747578 |
| GPR84 | 2.01809105 | 0.95403193 | 4.69483655 | 2.63E-05 | 0.0012583 | 2.43473106 |
| MYOF | 2.01584467 | 4.76808245 | 4.7990805 | 1.87E-05 | 0.00100277 | 2.69220592 |
| HERC6 | 2.00908938 | 6.1327843 | 4.90685962 | 1.31E-05 | 7.86E-04 | 3.01214841 |
| DHX58 | 2.00378908 | 5.59355218 | 3.89158561 | 3.34E-04 | 0.00697906 | -0.0585073 |
| SAMD4A | 2.00215697 | 4.46504118 | 4.23424064 | 1.15E-04 | 0.00338337 | 0.99344984 |

**Supplemental Table 8: Day 14 downregulated DEGs**

| **geneID** | **logFC** | **AveExpr** | **t** | **P.Value** | **adj.P.Val** | **B** |
| --- | --- | --- | --- | --- | --- | --- |
| ADAMTS5 | -4.2703246 | -0.0678239 | -5.2398812 | 4.37E-06 | 3.68E-04 | 2.96553106 |
| PHF24 | -4.1281247 | -0.4204843 | -4.5273428 | 4.52E-05 | 0.00175724 | 1.07075856 |
| RBM17P1 | -3.6292412 | 0.45420766 | -3.7529218 | 5.09E-04 | 0.00910743 | -0.3592191 |
| MTCYBP11 | -3.1386321 | 0.41697961 | -3.7791348 | 4.70E-04 | 0.00869732 | -0.2378834 |
| ENTPD2 | -3.0014965 | -1.0204361 | -5.2609943 | 4.07E-06 | 3.56E-04 | 2.73274732 |
| RP11-435F17.3 | -2.8793972 | 0.17552539 | -5.3132781 | 3.42E-06 | 3.21E-04 | 3.54846843 |
| OTX1 | -2.7834091 | 1.30226967 | -5.3924835 | 2.62E-06 | 2.82E-04 | 4.33858156 |
| ALPL | -2.7713125 | 6.55037767 | -3.8102068 | 4.28E-04 | 0.00821479 | -0.2659793 |
| RSPH14 | -2.7554649 | -0.178196 | -4.0774696 | 1.88E-04 | 0.00472545 | 0.41765881 |
| SCRT2 | -2.6695547 | 3.08530385 | -7.3037573 | 4.15E-09 | 5.45E-06 | 10.5645698 |
| TGM3 | -2.6673512 | 1.30725345 | -3.7247771 | 5.54E-04 | 0.00948169 | -0.2547174 |
| FABP6 | -2.6555066 | -0.0053424 | -3.8588684 | 3.69E-04 | 0.0074414 | -0.1282192 |
| TSPEAR | -2.6398871 | 0.73919599 | -4.4436994 | 5.92E-05 | 0.00211914 | 1.48180339 |
| CXCL6 | -2.4311896 | -0.6719037 | -4.6829926 | 2.73E-05 | 0.00127843 | 1.93745131 |
| RAB36 | -2.3856861 | 3.16647113 | -5.9520915 | 3.99E-07 | 9.62E-05 | 6.3853719 |
| RP4-673D20.4 | -2.3473481 | -1.1834619 | -4.4931758 | 5.05E-05 | 0.0019023 | 0.99557874 |
| AGMO | -2.336297 | 1.01712184 | -4.1931112 | 1.31E-04 | 0.00366963 | 0.96538114 |
| PDZD3 | -2.3339129 | 0.19178061 | -5.3585221 | 2.94E-06 | 2.97E-04 | 3.94552835 |
| RP11-34P1.2 | -2.2909288 | -0.8387437 | -5.0517082 | 8.15E-06 | 5.68E-04 | 2.40091969 |
| SDC2 | -2.2333407 | 0.18924219 | -4.0618828 | 1.98E-04 | 0.00482498 | 0.50261328 |
| THBD | -2.2104509 | 4.35362418 | -6.4547159 | 7.28E-08 | 3.53E-05 | 8.01732434 |
| CNTNAP3B | -2.1842551 | 4.25065745 | -3.7345248 | 5.38E-04 | 0.00931394 | -0.4111129 |
| CNTNAP3 | -2.14251 | 4.98229345 | -4.0936799 | 1.79E-04 | 0.00457207 | 0.57497861 |
| INHBB | -2.1030845 | -0.7925172 | -4.3931222 | 6.96E-05 | 0.00237789 | 0.9837876 |
| ARHGEF40 | -2.0702104 | 6.61822907 | -6.620432 | 4.15E-08 | 2.67E-05 | 8.55084276 |
| ANGPT1 | -2.0062407 | 1.99811146 | -4.444016 | 5.91E-05 | 0.00211914 | 1.74593849 |

**Supplemental Table 9: Overlap in DEGs on days 10 and 14**

|  | **Number** | **Genes** |
| --- | --- | --- |
| Unique Day 10 | 52 | AC003080.4, AC007899.3, AC007919.18, AC074338.4, ACO1, AIM2, BST2, CCR1, CD274, DDX58, DDX60L, EPHB2, EXOC3L1, HELZ2, IFIH1, IFIT5, IL1RN, KIAA1958, LAMP3, LGALS9, LIPA, LLpac-136A2.1, MS4A4A, MTND4P26, NUDT19P5, PARP12, PARP14, PARP9, PLSCR2, PML, PNPT1, RNA5SP39, RP11-466G12.3, RP11-820K3.2, SAMD9, SAMD9L, STAT2, TDRD7, TFEC, TNFAIP6, TNFSF10, TNFSF13B, TOR1B, TREX1, TRIM22, TRIM6, TTC21A, TTC26, USP41, XXbac-BPG541D20.6, ZBP1, ZCCHC2 |
| Shared Days 10 &14 | 60 | ADAMTS5, ANKRD22, ATF3, BATF2, C1QA, C1QB, C1QC, CARD17, CMPK2, CXCL6, DDX60, DHX58, EIF2AK2, EPSTI1, ETV7, FBXO39, FBXO6, FRMD3, GBP1, GBP4, HERC5, HERC6, IFI35, IFI44, IFI44L, IFI6, IFIT1, IFIT2, IFIT3, IFITM3, IRF7, LAP3, LY6E, MT2A, MX1, OAS1, OAS2, OAS3, OASL, OR52M2P, PLSCR1, RMI2, RP4-641G12.3, RPL37P6, RSAD2, RTP4, RUFY4, SAMD4A, SCO2, SERPING1, SIGLEC1, SMTNL1, SPATS2L, STAT1, TCN2, TMPRSS2, UBE2L6, USP18, XAF1, ZDHHC4P1 |
| Unique Day 14 | 117 | AGMO, ALPL, ANGPT1, ANLN, ANO5, ANXA10, ARHGEF40, ASPM, AURKB, BIRC5, BUB1, CCNB2, CDC25A, CDC45, CDCA2, CDCA5, CDK1, CENPA, CEP55, CNTNAP3, CNTNAP3B, CXCL11, CXCL9, CXCR2P1, DTL, E2F1, E2F8, ELOVL3, ENTPD2, FABP6, FAM72B, GBP1P1, GBP5, GPR84, HIST1H1B, HIST1H2AB, HIST1H2AH, HIST1H2AJ, HIST1H2AL, HIST1H2BE, HIST1H2BI, HIST1H2BL, HIST1H2BM, HIST1H2BO, HIST1H3B, HIST1H3C, HIST1H3F, HIST1H3G, HIST1H3J, HIST2H3A, HIST2H3C, HMMR, HNRNPCP4, IFI27, IGHV2-5, IGHV4-39, IGHV5-51, IGKV1D-16, IGKV2-28, IGLC1, IGLC2, IGLV1-40, IGLV1-44, IGLV1-47, IGLV1-51, IGLV2-23, IGLV2-8, IGLV3-1, IGLV3-21, IGLV3-25, IGLV6-57, IGLV8-61, INHBB, JCHAIN, KCTD14, KIF4A, KIFC1, LIF, MCM10, MKI67, MT1E, MTCYBP11, MYBL2, MYOF, MZB1, NAT8, NCAPG, NEURL3, NMRAL1P1, NUSAP1, OTX1, PBK, PDCD1LG2, PDZD3, PHF24, PXT1, PYCR1, RAB36, RBM17P1, RNASE1, RP11-34P1.2, RP11-435F17.3, RP4-673D20.4, RRM2, RSPH14, SCRT2, SDC1, SDC2, SF3A3P2, SOCS1, TGM3, THBD, TMEM51, TOP2A, TPX2, TSPEAR, TYMS |

**Supplemental Table 10: Components of Day 14 DEG modules**

| **Module 1** | ANLN, ASPM, AURKB, BIRC5, BUB1, CCNB2, CDC25A, CDC45, CDCA2, CDCA5, CDK1, CENPA, CEP55, CXCL9, DTL, E2F1, E2F8, HIST1H1B, HIST1H2AB, HIST1H2AH, HIST1H2AJ, HIST1H2AL, HIST1H2BE, HIST1H2BI, HIST1H2BL, HIST1H2BM, HIST1H2BO, HIST1H3B, HIST1H3C, HIST1H3F, HIST1H3G, HIST1H3J, HIST2H3A, HIST2H3C, HMMR, IGHV2-5, IGHV4-39, IGHV5-51, IGKV1D-16, IGKV2-28, IGLC1, IGLC2, IGLV1-40, IGLV1-44, IGLV1-47, IGLV1-51, IGLV2-23, IGLV2-8, IGLV3-1, IGLV3-21, IGLV3-25, IGLV6-57, IGLV8-61, JCHAIN, KIF4A, KIFC1, MCM10, MKI67, MYBL2, MZB1, NCAPG, NUSAP1, PBK, PYCR1, RRM2, SDC1, TOP2A, TPX2, TYMS |
| --- | --- |
| **Module 2** | ANKRD22, ANO5, ANXA10, ATF3, BATF2, C1QA, C1QB, C1QC, CARD17, CMPK2, CXCL11, CXCR2P1, DDX60, DHX58, EIF2AK2, ELOVL3, EPSTI1, ETV7, FAM72B, FBXO39, FBXO6, FRMD3, GBP1, GBP1P1, GBP4, GBP5, GPR84, HERC5, HERC6, HNRNPCP4, IFI27, IFI35, IFI44, IFI44L, IFI6, IFIT1, IFIT2, IFIT3, IFITM3, IRF7, KCTD14, LAP3, LIF, LY6E, MT1E, MT2A, MX1, MYOF, NAT8, NEURL3, NMRAL1P1, OAS1, OAS2, OAS3, OASL, OR52M2P, PDCD1LG2, PLSCR1, PXT1, RMI2, RNASE1, RP4-641G12.3, RPL37P6, RSAD2, RTP4, RUFY4, SAMD4A, SCO2, SERPING1, SF3A3P2, SIGLEC1, SMTNL1, SOCS1, SPATS2L, STAT1, TCN2, TMEM51, TMPRSS2, UBE2L6, USP18, XAF1, ZDHHC4P1 |
| **Module 3** | ADAMTS5, AGMO, ALPL, ANGPT1, ARHGEF40, CNTNAP3, CNTNAP3B, CXCL6, ENTPD2, FABP6, INHBB, MTCYBP11, OTX1, PDZD3, PHF24, RAB36, RBM17P1, RP11-34P1.2, RP11-435F17.3, RP4-673D20.4, RSPH14, SCRT2, SDC2, TGM3, THBD, TSPEAR |
