## Supplemental figures for "Evolution of inflammation and immunity in a dengue virus 1 human infection model"

A)

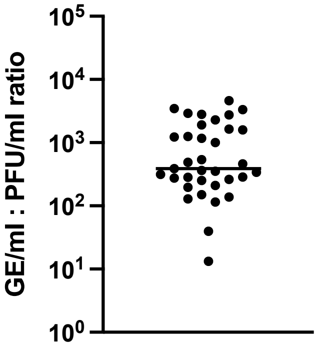


**Supplemental figure 1. Kinetics of DENV-1 viremia as assessed by Vero plaque assay.** Ratio of viral RNA (GE/ml) to infectious virus (PFU/ml) in serum from all time points with > 100 PFU/ml

A)

MNSRSTSLSMTCIAVGMVTLYLGVMVQADSGCVINWKGRELKCGSGIFVTNEVHTWTEQYKFQADSPKRLSAAIGK
AWEEGVCGIRSATRLENIMWKQISNELNHILLENDMKFTVVVGDVSGILAQGKKMIRPQPMEHKYSWKSWGKAKII
GADVQNTTFIIDGPNTPECPDNQRAWNIWEVEDYGFGIFTTNIWLKLRDSYTQVCDHRLMSAAIKDSKAVHADMGY
WIESEKNETWKLARASFIEVKTCIWPKSHTLWSNGVLESEMIIPKIYGGPISQHNYRPGYFTQTAGPWHLGKLELD
FDLCEGTTVVVDEHCGNRGPSLRTTTVTGKTIHEWCCRSCTLPPLRFKGEDGCWYGMEIRPVKEKEENLVKSMVSA
GSGEVDSFSLGLLCISIMIEEVMRSRWSRKMLMTGTLAVFLLLTMGQLTWNDLIRLCIMVGANASDKMGMGTTYLA
LMATFRMRPMFAVGLLFRRLTSREVLLLTVGLSLVASVELPNSLEELGDGLAMGIMMLKLLTDFQSHQLWATLLSL
TFVKTTFSLHYAWKTMAMILSIVSLFPLCLSTTSQKTTWLPVLLGSLGCKPLTMFLITENKIWGRK

B)

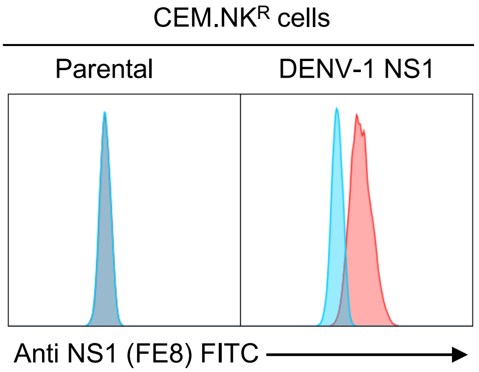


C)

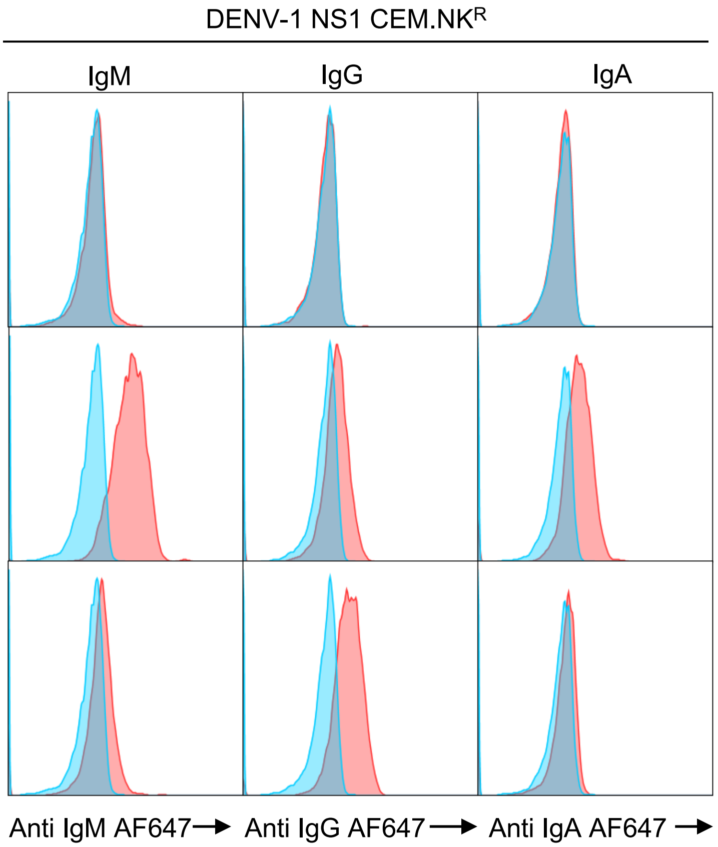


**Supplemental figure 2.** NS1 opsonization assay. **A) S**equence and annotation of the DENV-1 NS1 expression construct used in this study. Yellow = signal peptide, green = NS1, blue = NS2A. Sequence derived from WestPac74 (U88535.1) **B)** Anti-NS1 staining mAb staining (clone FE8) of parental CEM.NKR cell line and DENV-1 NS1 expressing CEM.NKR cells. Unstained cells shown in blue, FE8 stained shown in red **C)** Representative NS1 opsonizing activity of serum collected at days 0, 24, and 90 days post DENV-1 challenge. Unstained cells shown in blue, serum stained shown in red

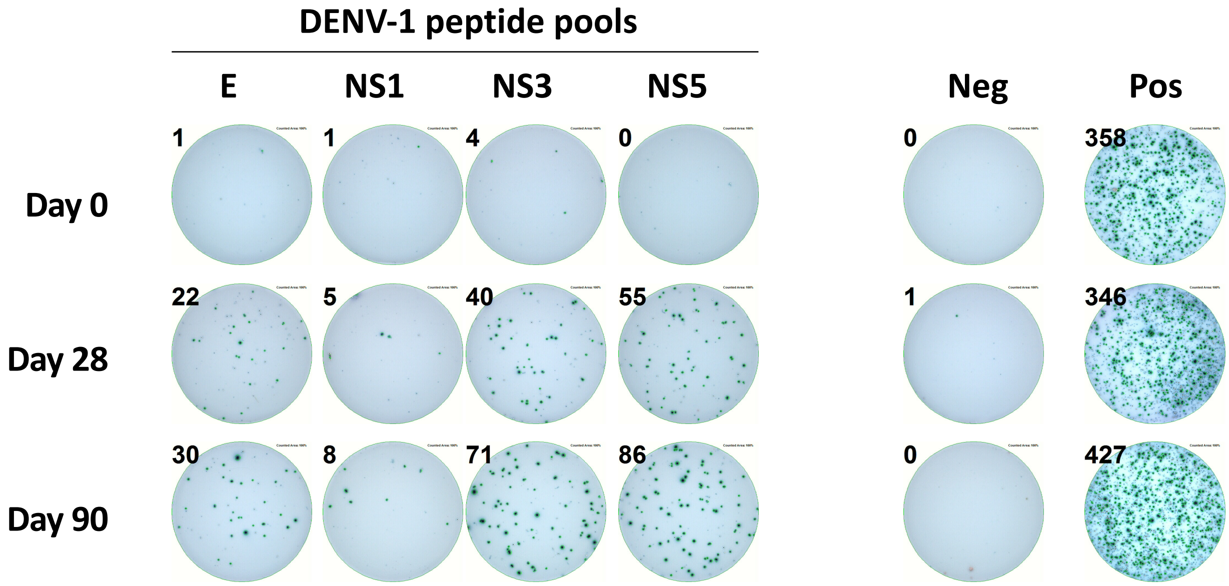


**Supplemental figure 3.** Representative IFN-γ ELISPOT assay images from samples obtained 0, 28 and 90 days post DENV-1 infection.
